# Developmental embedding of parvalbumin interneurons drives local and crosshemispheric prefrontal gamma synchrony

**DOI:** 10.1101/2025.05.12.653375

**Authors:** Anton Offermanns, Jastyn A. Pöpplau, Ileana L. Hanganu-Opatz

## Abstract

Gamma oscillations are a pivotal trait of cortical cognitive processing. However, the ability to generate gamma oscillations evolves with age and requires cellular adjustments of the underlying neural networks. In the prefrontal cortex, gamma oscillations emerge relatively late compared to other cortical areas, yet the developmental mechanisms leading to the generation of adult-like gamma oscillations are poorly understood. Here, we combine bilateral *in vivo* electrophysiology and selective optogenetic manipulations of parvalbumin- (PV+) and somatostatin-positive (SOM+) interneurons in the mouse medial prefrontal cortex along late development to investigate their role for the age-dependent maturation of gamma oscillations. We show that crosshemispheric gamma synchrony strengthens with age, in line with the previously reported increase in local gamma power. Following a similar timeline, the inhibitory effect of PV+ interneurons emerges which start to functionally operate within the classical gamma frequency range from adolescence onwards. In contrast, SOM+ interneurons show no such age-dependent functional integration and display their beta oscillation modulating inhibitory effect across age. These data identify the SOM+ to PV+ interneuron switch as a mechanism of gamma ontogeny and emergence of crosshemispheric synchrony in the developing prefrontal cortex.

**Graphical abstract:** 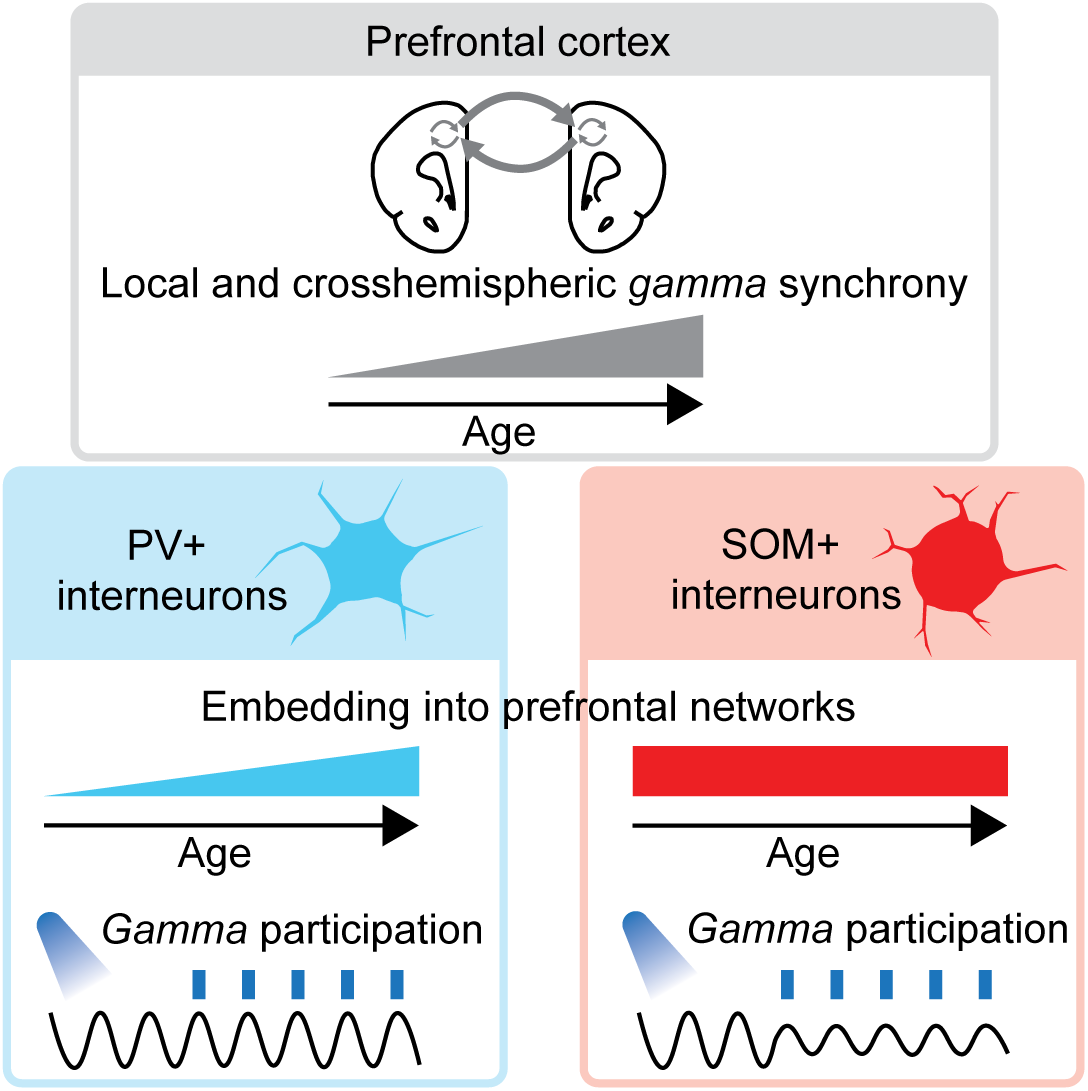

## Introduction

The prefrontal cortex (PFC), which can be considered the brain’s core processor, generates oscillation in gamma frequency range (broadly defined as 30-100 Hz) to solve complex cognitive tasks. ^1–3^ The local and crosshemispheric synchronization of gamma rhythms has been causally linked to attention, decision-making, and learning. ^4–8^ On the flip side, both local and crosshemispheric gamma synchrony ^9–15^ as well as cognitive functions ^16^ are impaired in several neurological and neuropsychiatric disorders. Moreover, transient abnormal oscillatory patterns have been related to the pathogenesis of such diseases and are potentially caused by an early imbalance of the GABAergic system. ^4,10,13,17^ Consequently, disentangling the physiological developmental processes that lead to the generation of rhythmic oscillatory activity is of particular relevance for understanding the pathomechanisms of impaired brain development.

Adult-like fast (>30 Hz) oscillations emerge during the second postnatal week in the hippocampus, visual, and somatosensory cortices of mice. ^18–22^ In contrast, fast oscillations in the mouse medial PFC (mPFC) show a protracted development, increasing in power and frequency along second to third postnatal week. At the end of the fourth postnatal week, the acceleration of oscillations settles at adult-like frequencies between 50 and 60 Hz. ^23–25^ This initial shift from beta/slow gamma to faster gamma oscillations is followed by a transient peri-adolescent decrease in oscillatory power but not frequency. ^25^ Notably, the protracted emergence of mature gamma oscillations in the PFC aligns with the delayed emergence of cognitive functions and relates to the onset of neuropsychiatric symptoms in mental diseases. ^26–28^ In primary sensory cortices, the emergence of gamma oscillations has been attributed to functional shifts in the GABAergic interneuron (IN) population. ^20,29–35^ For example, while after birth, inhibition is mainly exerted by somatostatin-positive (SOM+) INs, during the first two postnatal weeks a switch towards a stronger inhibition mediated by fast-spiking (FS) INs, putatively parvalbumin-positive (PV+) ^36^, takes place. This switch possibly allows for more efficient processing of sensory information. ^30,31^

In the PFC, PV+ and SOM+ INs are the two most abundant subpopulations of GABAergic INs and take on different roles in cortical computations at adult age. ^37–43^ SOM+ INs inhibit other neurons at the dendritic tree, whereas PV+ INs target the soma, allowing for faster and more precisely timed inhibition. ^40^ Not only their distinct connectivity, but also faster intrinsic kinetic properties, in comparison to other INs, make PV+ INs particularly fast signaling units. ^36,40^ This most likely results in SOM+ INs participating mainly in the generation of beta oscillations, ^44,45^ while PV+ INs are suited to form microcircuits that generate gamma oscillations. ^4,5,36,44,46,47^ Additionally, callosal projections of prefrontal PV+ INs have been found to control crosshemispheric gamma synchrony during cognitive tasks in adult mice. ^6,48^ In parallel to the developmental increase in power and frequency of prefrontal gamma oscillations, PV+ IN physiology matures and parvalbumin expression increases along late development, whereas SOM+ INs do not show such age-related changes during late development. ^23,35,49–55^ The developmental dynamics of PV+ INs particularly affect layer 2/3 (L2/3), ^35,51,53^ which we previously identified as origin of gamma oscillation generation across development. ^24,56^ Therefore, it is likely that a developmental embedding of PV+ INs in prefrontal networks contributes to the protracted maturation of gamma oscillations and crosshemispheric synchrony in the mPFC. ^57^

Here, we aim to test this hypothesis and address the cellular mechanisms of local as well as crosshemispheric gamma development, by combining bilateral extracellular recordings with optogenetic manipulation of PV+ and SOM+ INs in the mPFC of non-anesthetized postnatal day (P)16-60 mice. For this we assessed (i) the developmental dynamics of crosshemispheric prefrontal gamma synchrony, (ii) the age-dependent inhibitory function of PV+ and SOM+ INs in prefrontal networks, and (iii) how PV+ and SOM+ INs contribute to the developmental dynamics of local and crosshemispheric gamma synchrony in the mPFC. We provide evidence for an age-dependent functional integration of PV+, but not SOM+, INs in prefrontal networks which controls the emergence of local gamma power and crosshemispheric synchrony.

## Results

### Crosshemispheric gamma synchrony increases over age and is driven by prefrontal L2/3

To investigate developmental changes in local and crosshemispheric communication in the prelimbic subdivision (PL) of the mouse mPFC, we gathered a dataset of simultaneously recorded local field potential (LFP) and single-unit activity (SUA) from both hemispheres. For this, we used PV-Cre^+^, SOM-Cre^+^, and wildtype (WT) mice between P16 and P60 (divided into four age groups: P16-17, P20-21, P30-33, P50-60). The mice were chronically implanted with a head plate and recorded in a non-anesthetized state on the Mobile HomeCage. To manipulate and monitor the prefrontal activity patterns as well as local and crosshemispheric communication in a layer-specific manner, we acutely inserted a four-shank optrode, spanning across all layers, in the right and a one-shank optrode in the left mPFC (Fig. 1A). To account for random-effects, such as sex, statistical analysis was conducted using linear mixed-effect models (see Table S1 for details). As previously reported ^23^, the broad-band oscillatory power and neuronal spiking augmented with age and showed wide temporal similarity of oscillatory patterns across layers and hemispheres (Fig. 1B).

**Figure 1.**
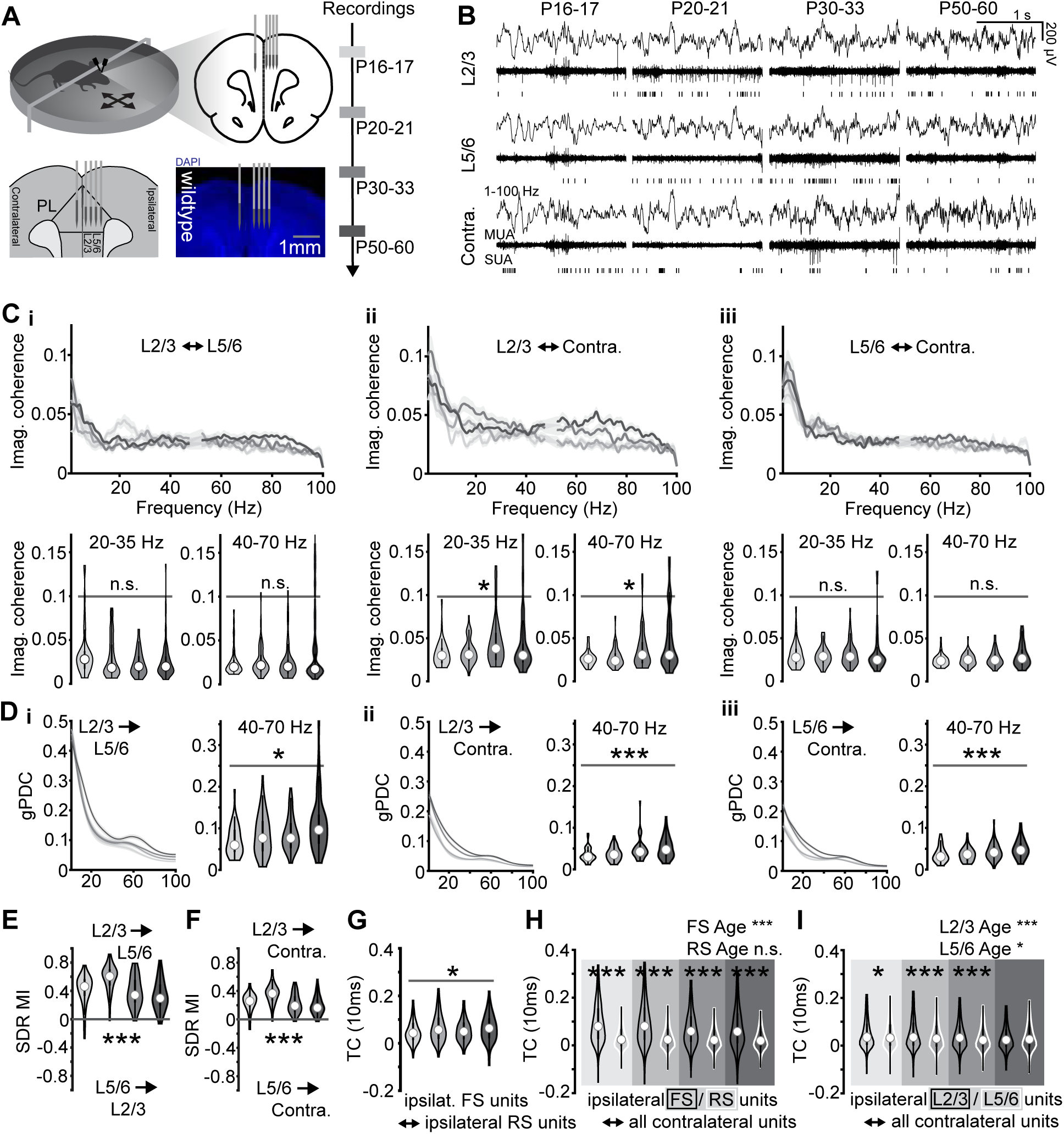
Local and crosshemispheric mPFC communication throughout development. (**A**) Left, schematic and digital photomontage of mouse head-fixation, 4-shank optrode positioning in the right mPFC hemisphere (ipsilateral), and 1-shank optrode positioning in the left mPFC hemisphere (contralateral). Right, timeline and age-dependent colorcodes of recordings. (**B**) Simultaneosly recorded 1-100 Hz filtered LFP, 500-9000 Hz filtered LFP (multi-unit activity, MUA) and SUA in L2/3, L5/6 of the ipsilateral as well as contralateral mPFC for each age group. (**C i**) Top, averaged imaginary coherence between simultaneously recorded LFP in ispsilateral L2/3 and L5/6 of the mPFC for each age group (n=236 recordings, 110 mice). Bottom left, violinplots displaying imaginary coherence between 20 and 35 Hz for each age group. Bottom right, violinplots displaying imaginary coherence between 40 and 70 Hz for each age group. Due to possible electrical artifacts in the 50 Hz range, values between 47 and 53 Hz were omitted for visualization and quantification. (**C ii**) Same as (C i) for imaginary coherence between ipsilateral L2/3 and contralateral mPFC. (**C iii**) Same as (C i) for imaginary coherence between ipsilateral L5/6 and contralateral mPFC. (**D i**) Left, averaged gPDC indicating the information flow from ipsilateral L2/3 to ipsilateral L5/6 of the mPFC for each age group (n=234 recordings, 111 mice). Right, violinplots displaying the gPDC between 40-70 Hz for each age group. (**D ii**) Same as (D i) for information flow from ispsilateral L2/3 to contralateral mPFC. (D iii) Same as (D i) for information flow from ispsilateral L5/6 to contralateral mPFC. (**E**) Violinplots displaying the MI of the SDR between L2/3 -> L5/6 and L5/6 -> L2/3 in the ipsilateral mPFC for each age group (n=233 recordings, 110 mice). (**F**) Violinplots displaying the MI of the SDR between ipsilateral L2/3 -> contralateral mPFC and between ipsilateral L5/6 -> contralateral mPFC for each age group (n=233 recordings, 110 mice). (**G**) Violinplots displaying the TC between simultaneously recorded spike trains of ipsilateral FS units and ipsilateral RS units for each age group (n=5216 unit pairs, 211 recordings, 108 mice). Time lag for TC calculation = 10ms. (**H**) Violinplots displaying the TC between simultaneously recorded spike trains of ipsilateral FS (black outline, n=7420 unit pairs, 189 recordings, 102 mice) and RS (white outline, n=24489 unit pairs, 205 recordings, 105 mice) units and contralateral units for each age group. Time lag for TC calculation = 10ms. Asterisks in the plot indicate significant differences between FS <> contralateral and RS <> contralateral unit pairs for all age groups. Asterisks in the upper right corner indicate a significant age effect. *** p < 0.001, linear mixed-effect models. (**I**) Violinplots displaying the TC between simultaneously recorded spike trains of ipsilateral L2/3 (black outline, n=15337 unit pairs, 196 recordings, 102 mice) or L5/6 (white outline, n=16510 unit pairs, 201 recordings, 105 mice) units and contralateral units for each age group. Time lag for TC calculation = 10ms. Asterisks in the plot indicate significant difference between L2/3 <> contralateral and L2/3 <> contralateral unit pairs for all age groups. Asterisks in the upper right corner indicate significant age effect. * p < 0.05, *** p < 0.001, linear mixed-effect models. Lineplots in (C + D) represent the mean ± standard error of the mean per age group. Violinplots in (C)-(I) represent the median with 25^th^ and 75^th^ percentile. Asterisks in (C)– (G) indicate significant effect of age. * p < 0.05, ** p < 0.01, *** p < 0.001, linear mixed-effect models. See Table S1 for detailed statistics.

First, we analyzed layer-specific prefrontal network activity to resolve changes in local and crosshemispheric communication along development (Fig. 1C). For this, we calculated the imaginary part of the coherence, which is insensitive to false connectivity arising from volume conduction. ^58^ We found that beta (20-35 Hz) coherence between L2/3 and the contralateral mPFC increases until P30-33, whereas gamma (40-70 Hz) coherence increases until adulthood. In contrast, the local coherence between L2/3 and L5/6, as well as coherence between L5/6 and the contralateral mPFC did not significantly change with age (Fig. 1C i-iii). Strengthening of gamma synchrony is driven by L2/3 as revealed by quantifying the generalized partial directed coherence (gPDC) ^59^ which progressively increased with age, both within and between hemispheres (Fig. 1 D). Similarly, as indicated by a positive modulation index (MI) of the spectral density ratio (SDR) ^60,61^, L2/3 activity drives local and crosshemispheric coupling across all frequencies more efficiently than L5/6 (Fig. 1E, F). This effect peaks around P20 and decreases as the mice reach adulthood, paralleling the onset of fast activity in the mPFC. ^23^

Second, we investigated the age- and layer-dependent dynamics of spiking activity to get first insights into the neuronal basis of local and crosshemispheric communication in the developing mPFC. For this, we calculated the tiling coefficient (TC) of SUA to assess spike time correlations that are unbiased by firing rates. ^62^ We classified all units according to their extracellular waveform shape as FS (i.e. putative PV+ INs) and regular-spiking (RS) (i.e. mainly putative pyramidal neurons) units ^23,36^ as well as according to their anatomical position in L2/3 or L5/6. We calculated the TC with a 10 ms lag as a proxy of functional mono/disynaptic excitatory connections between all locally and crosshemispherically recorded unit pairs. In the same hemisphere, the correlation of firing between FS-RS unit pairs significantly increased with age (Fig. 1G), whereas the correlations between all other unit pairs were constant along development (Fig. S1A i-iii). Solely, the correlation between RS-RS unit pairs, despite being weaker compared to FS-FS pairs, slightly peaked around P30 (Fig. S1A iv). When analyzing the prefrontal firing across hemispheres, we found that throughout development FS units were more correlated with units in the contralateral mPFC than RS units (Fig. 1H). Similarly, units in L2/3 fire more correlated with contralateral units than L5/6, yet the TC values were comparable at adult age (Fig. 1I). Generally, the crosshemispheric firing decorrelated with age, which reached significance for FS units across layers and RS units in L2/3, but not in L5/6 (Fig. S1B).

Thus, the developmental increase in local gamma power and spike time correlations between FS and RS units is accompanied by a strengthening of crosshemispheric gamma synchrony and decorrelation of SUA. Moreover, these results highlight L2/3 as driver of age-dependent changes in prefrontal network activity.

PV+ but not SOM+ interneuron-mediated inhibition in local prefrontal networks strengthens with age To gain a more detailed understanding of the cellular interactions underlying the above-described age-dependent changes in local and crosshemispheric coupling within prefrontal networks, we investigated the local inhibitory function of PV+ and SOM+ INs over age. For this, PV-Cre^+^, SOM-Cre^+^, and WT mice were injected with a Cre-dependent virus, encoding for channelrhodopsin 2, into the right mPFC hemisphere. PV+ or SOM+ INs were activated in a layer specific manner by light (3 ms pulses, 473 nm, 30 Hz train), while recording the neuronal firing across all layers (Fig. 2A). Since no significant differences were obtained after light stimulation of INs in L2/3 and L5/6 (Fig. S2A), data from all layers were pooled.

**Figure 2.**
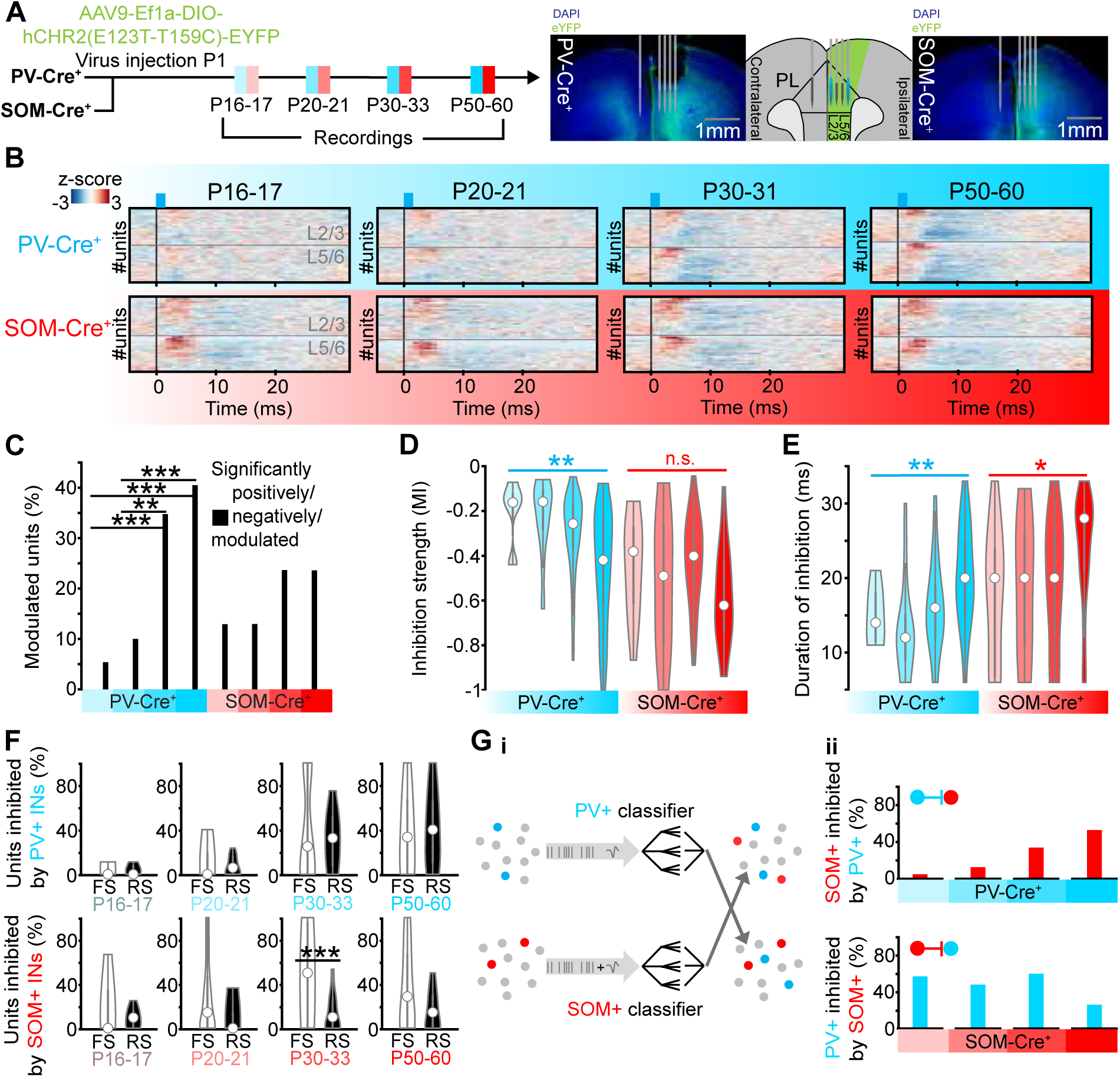
Activation of PV+ and SOM+ interneurons throughout development. (**A**) Left, timeline as well as colorcode of injection and recording ages of PV-Cre^+^ and SOM-Cre^+^ mice. Right, schematic drawing of virus expression (green area), optrode positions of 4-shank optrode in right mPFC (ipsilateral) and 1-shank optrode in left mPFC (contralateral), and sites of optogenetic stimulation (blue beams). Digital photomontages showing the optrode positions and virus expression in a coronal slice of a P60 PV-Cre and P60 SOM-Cre^+^ mouse. (**B**) Rasterplots showing the z-scored and stimulation period averaged spike trains of all recorded units in PV-Cre^+^ and SOM-Cre^+^ mice for each age group during optogenetic stimulation with blue (473 nm wavelength) 3 ms-long light pulses delivered in the ipsilateral mPFC with a frequency of 30 Hz. Units recorded in L2/3 are shown above and units recorded in L5/6 are shown below the grey horizontal lines. (PV-Cre^+^ mice: P16-17 n=316 units, P20-21 n=478 units, P30-33 n=361 units, P50-60 n=442 units; SOM-Cre^+^ mice: P16-17 n=322 units, P20-21 n=300 units, P30-33 n=277 units, P50-60 n=268 units) (**C**) Barplots displaying the absolute percentage of significantly modulated units after optogenetic 30 Hz pulse stimulation in L2/3 or L5/6 in PV-Cre^+^ (n=110 recordings, 53 mice) and SOM-Cre^+^ (n=119 recordings, 52 mice) mice for each age group. Individual percentages were not calculated since binomial models were used for statistical analysis. Asterisks indicate significant difference between age groups. ** p < 0.01, *** p < 0.001, generalized linear mixed-effect models. (**D**) Violinplots displaying the inhibition strength of significantly negatively modulated units after optogenetic pulse stimulation with a frequency of 30 Hz in PV-Cre^+^ (blue, n=289 units, 110 recordings, 53 mice) or SOM-Cre^+^ (red, n=149 units, 119 recordings, 52 mice) mice for each age group. Asterisks indicate significant effect of age. ** p < 0.01, linear mixed-effect models. (**E**) Violinplots displaying the duration of inhibition, measured as the time units need to reach their pre pulse firing rate, of significantly negatively modulated units after optogenetic pulse stimulation with a frequency of 30 Hz in PV-Cre^+^ (blue, n=289 units, 110 recordings, 53 mice) or SOM-Cre^+^ (red, n=149 units, 119 recordings, 52 mice) mice for each age group. Asterisks indicate significant effect of age. * p < 0.05, ** p < 0.01, linear mixed-effect models. (**F**) Violinplots displaying the percentage of significantly negatively modulated FS and RS units per animal after optogenetic pulse stimulation with a frequency of 30 Hz in PV-Cre^+^ (n=107 recordings, 53 mice) or SOM-Cre^+^ (n=106 recordings, 50 mice) mice for each age group. Asterisks indicate significant difference between the percentage of negatively modulated FS and RS units for all age groups. *** p < 0.001, linear mixed-effect models. (**G i**) Schematic drawing of PV+/SOM+ classification procedure. (**G ii**) Top, barplots displaying the percentage of significantly negatively modulated putative SOM+ units (based on classifier results) after optogenetic pulse stimulation in PV-Cre^+^ (n=110 recordings, 53 mice) mice for each age group. Bottom, barplots displaying the percentage of significantly negatively modulated putative PV+ units (based on classifier results) after optogenetic pulse stimulation in SOM-Cre^+^ (n=119 recordings, 52 mice) mice for each age group. Violinplots in (D)-(F) represent the median with 25^th^ and 75^th^ percentile. See Table S1 for detailed statistics.

Activation of PV+ and SOM+ INs had different effects on prefrontal firing along development. The fraction of significantly activated and the fraction of inhibited units after activation of PV+ INs, but not SOM+ INs, increased with age, although the age-dependent increase in positive modulation after activation of PV+ INs (linear mixed-effect model, Chi-Square: 12.922, p = 0.0048) did not reach significance in post-hoc analyses (Fig. 2B, C; Table S1). Similarly, the strength of inhibition, which was assessed as MI of optogenetic stimulation (i.e. lower the values, stronger the inhibition; see Methods), exerted by PV+, but not by SOM+, INs augmented with age. However, the overall inhibition strength of SOM+ INs was stronger than that of PV+ INs (Fig. 2D), potentially as result of generally higher SOM+ IN densities in mPFC. ^63^ Moreover, the duration of inhibition after PV+ IN activation progressively increased over age, while it was rather constant for SOM+ INs along development with a minor increase towards adulthood (Fig. 2E). Mirroring the maturation of cellular properties, the FS and RS clusters became better separated with age, which occurred especially within L2/3 (Fig. S2D). These findings were not biased by changing proportions of FS and RS units, as FS units make up about 25% of all units independent of age during the investigated time window (Fig. S2E).

Next, we investigated, which cell types are inhibited by PV+ and SOM+ INs along development. At adult age, PV+ and SOM+ INs target not only PYRs but also each other. ^64^ First, to assess the developmental dynamics of these interactions, we clustered the units inhibited by PV+ and SOM+ INs into FS and RS units. While PV+ INs equally inhibited FS and RS units and no age-dependent changes were detected, SOM+ INs inhibited FS stronger than RS, with the strongest effects detected at P30-33 (Fig. 2F). Optogenetic photo-tagging of PV+ and SOM+ INs revealed that PV+ INs in the mPFC mainly consist of FS units, whereas SOM+ INs equally consist of FS and RS units (Fig. S2F). Thus, the stronger inhibition of FS units by SOM+ INs suggests that SOM+ INs form robust inhibitory connections with PV+ INs along development. Second, to gain more direct insights into age-dependent interactions between PV+-SOM+ INs, we used the full dataset of all recorded units, including photo-tagged PV+ and SOM+ INs, and trained two classifiers with the aim of detecting PV+ or SOM+ INs based on firing and waveform features. Running the PV+ classifier on the dataset of units recorded in SOM-Cre^+^ mice enabled us to detect classified putative PV+ INs and optogenetically photo-tagged SOM+ INs recorded in the same animal. Likewise, we ran the SOM+ classifier on the dataset of units recorded in PV-Cre^+^ mice (Fig. 2G; see Methods for a detailed description of the classifier). By these means, we investigated the reciprocal inhibitory effect of both IN types directly. We found that PV+ INs increasingly inhibit SOM+ INs with age (Fig. 2G ii). The portion of PV+ INs inhibited by SOM+ INs, despite being disproportionally higher compared with the portion of units inhibited by SOM+ INs (Fig. 2C), decreased toward adult age. Additionally, the observed changes were not affected by off-target effects since light stimulation in control WT mice led to no changes in firing activity (Fig. S2B, C).

Taken together, the results show that the inhibitory function of PV+ INs in prefrontal networks is developmentally protracted compared to SOM+ INs. Moreover, the mutual inhibition between PV+ and SOM+ INs shifts with age towards a stronger inhibition of PV+ INs onto SOM+ INs. PV+ but not SOM+ INs start to functionally operate within the gamma frequency range during adolescence To assess whether the developmental strengthening of PV+ IN-mediated inhibition contributes to the frequency acceleration from beta to gamma oscillations along development, we compared the effects of optogenetic pulse stimulation of PV+ and SOM+ INs at either 30 or 50 Hz. The number of activated PV+ and SOM+ INs was independent of the stimulation frequency (Fig. 3B i). However, SOM+ INs followed 30 Hz pulse stimulation better than 50 Hz stimulation, while PV+ INs followed both stimulation frequencies equally (Fig. 3B ii). Consequently, the portion of units inhibited by light-activation of SOM+, but not PV+ INs, was significantly smaller after 50 Hz stimulation when compared to 30 Hz stimulation in P30-33 and P50-60 mice (Fig. 3B iii). This suggests that PV+ IN-mediated inhibition is efficient over a broad band of fast frequencies (i.e. corresponding to beta and gamma bands), whereas SOM+ IN-mediated inhibition is only efficient in the lower frequency range (i.e. corresponding to beta band), presumably caused by slower kinetics and firing properties ^40^ of SOM+ INs.

**Figure 3.**
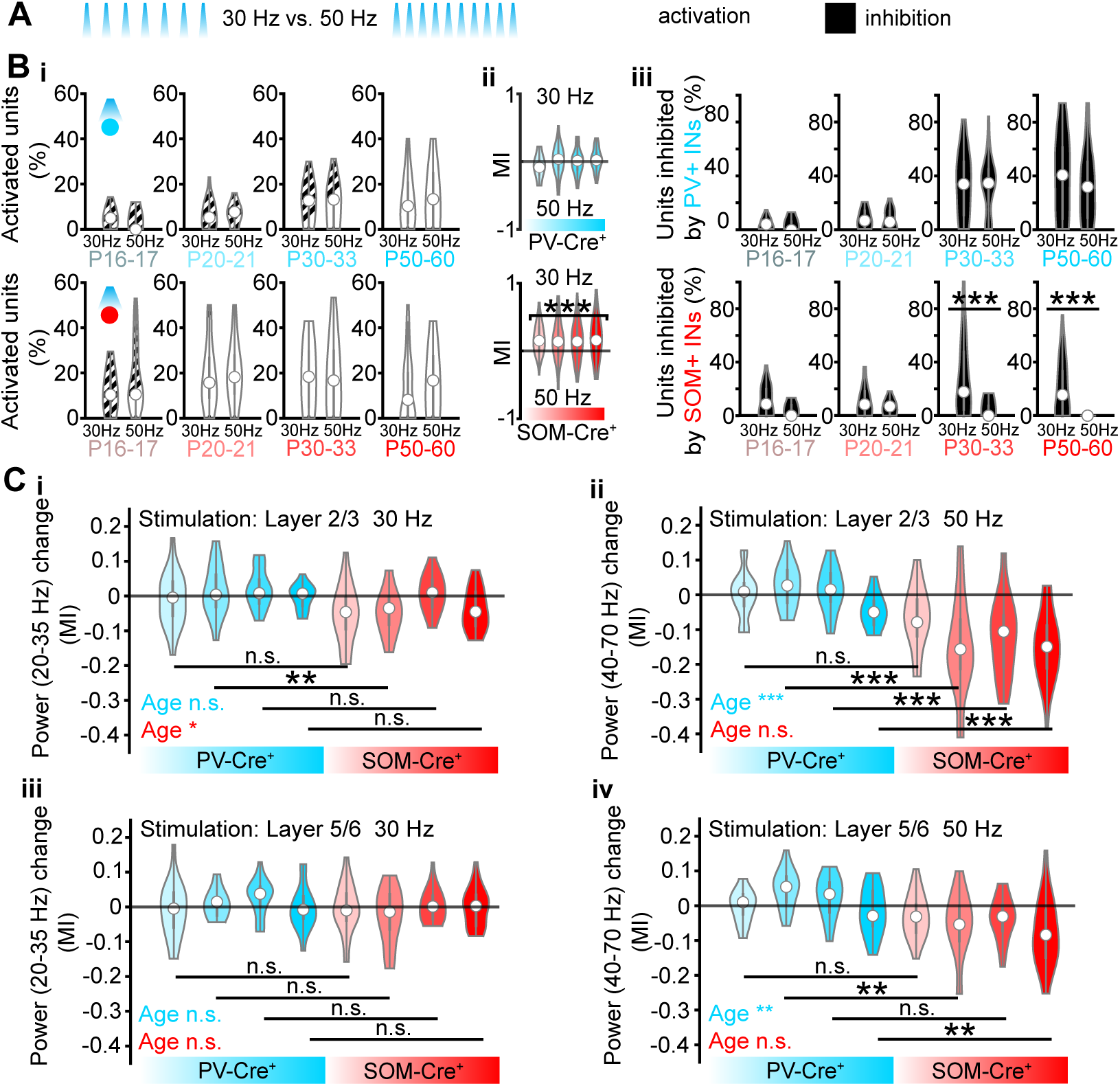
Effects of rhythmic stimulation of PV+ and SOM+ interneurons. (**A**) Schematic of optogenetic pulse stimulation with a frequency of 30 Hz and 50 Hz. (**B i**) Violinplots displaying the percentage of significantly positively modulated units per animal after optogenetic pulse stimulation in the mPFC of PV-Cre^+^ (top, n=106 recordings, 53 mice) and SOM-Cre^+^ (bottom, n=106 recordings, 50 mice) mice with 30 and 50 Hz for each age group. (**B ii**) Violinplots displaying the MI of the portion of pulses followed by a spike during 30 and 50 Hz optogenetic pulse stimulation in the mPFC of PV-Cre^+^ (top, n=106 recordings, 53 mice) and SOM-Cre^+^ (bottom, n=106 recordings, 50 mice) mice for each age group. Asterisk indicates significant difference between 30 Hz and 50 Hz stimulation for all age groups. *** p < 0.001, Wilcoxin signed-rank test. (**B iii**) Violinplots displaying the percentage of significantly negatively modulated units per animal after optogenetic pulse stimulation in the mPFC of PV-Cre^+^ (top, n=106 recordings, 52 mice) and SOM-Cre^+^ (bottom, n=106 recordings, 49 mice) mice with 30 and 50 Hz for each age group. Asterisks indicate significant difference between the percentage of negatively modulated units after 30 Hz and 50 Hz stimulation for all age groups. *** p < 0.001, linear mixed-effect models. (**C i+iii**) Violinplots displaying the MI of LFP power in the 20-35 Hz range between baseline recording and optogenetic 30 Hz pulse stimulation in L2/3(i) and L5/6 (iii) in the mPFC of PV-Cre^+^ (blue, n=102 recordings, 35 mice) and SOM-Cre^+^ (red, n=105 recordings, 36 mice) mice for each age group. (**C ii+iv**) Violinplots displaying the MI of LFP power in the 40-70 Hz range between baseline recording and optogenetic 50 Hz pulse stimulation in L2/3 (ii) and L5/6 (iv) in the mPFC of PV-Cre^+^ (blue, n=102 recordings, 35 mice) and SOM-Cre^+^ (red, n=105 recordings, 36 mice) mice for each age group. Violinplots in (B + C) represent the median with 25^th^ and 75^th^ percentile. In (B) a unit is considered being significantly positively modulated, if it is significantly positively modulated after optogenetic stimulation in L2/3 or L5/6. In (C) colored asterisks indicate a significant effect of age. Black asterisks indicate significant differences between stimulation of PV-Cre^+^ and SOM-Cre^+^ mice for all age groups. * p < 0.05, ** p < 0.01, *** p < 0.001, linear mixed-effect models. See Table S1 for detailed statistics.

Next, we tested whether the identified frequency specificity of PV+ and SOM+ IN-mediated inhibition is relevant for the local generation of fast oscillations. To this aim, we compared the impact of 30 and 50 Hz pulse stimulation of PV+ and SOM+ INs located in L2/3 and L5/6 on the power of beta and gamma oscillations. Stimulating PV+ or SOM+ INs with 30 Hz pulses did not substantially affect the generation of endogenous beta oscillations (Fig. 3C). However, using a stimulation frequency of 50 Hz disrupted the power of endogenous gamma oscillation when stimulating SOM+ but not PV+ INs, indicating that SOM+ INs are not mechanistically involved in the generation of gamma oscillations. 50 Hz stimulation of SOM+ INs might result in a slower or out-of-frequency inhibition of other neurons ^44^, leading to the disruption of endogenous gamma oscillations (Fig. 3B iii). This effect increased with age and is more prominent in L2/3 where gamma oscillations are generated. ^24,56^

Thus, both PV+ and SOM+ INs can be involved in the generation of beta oscillations but only PV+ INs can functionally operate in the gamma frequency range, suggesting a critical involvement of their embedding into prefrontal networks for the emergence of gamma oscillations along development.

### PV+ and SOM+ interneuron activation differently affects crosshemispheric communication

Finally, to assess how PV+ and SOM+ INs contribute to the crosshemispheric prefrontal communication along development, we compared the effects of their activation by 30 and 50 Hz light pulses on crosshemispheric oscillatory synchrony. We analyzed the crosshemispheric coherence in the beta and gamma frequency range upon stimulation in either L2/3 or L5/6. Light stimulation at 30 Hz of PV+ or SOM+ INs in L2/3 or L5/6 did not affect the endogenous crosshemispheric coherence in the beta range (Fig. 4A i+iii). In contrast, stimulation of SOM+ INs with 50 Hz light pulses disrupted the crosshemispheric endogenous gamma coherence increasingly with age which was most prominent in L2/3 (Fig. 4A ii+iv). This is similar to the effect of SOM+ IN stimulation on local gamma generation and indicates that PV+, but not SOM+ INs, are functionally involved in crosshemispheric gamma synchronization throughout development. Moreover, these results confirm the major contribution of L2/3 as driver of crosshemispheric prefrontal gamma synchrony.

**Figure 4.**
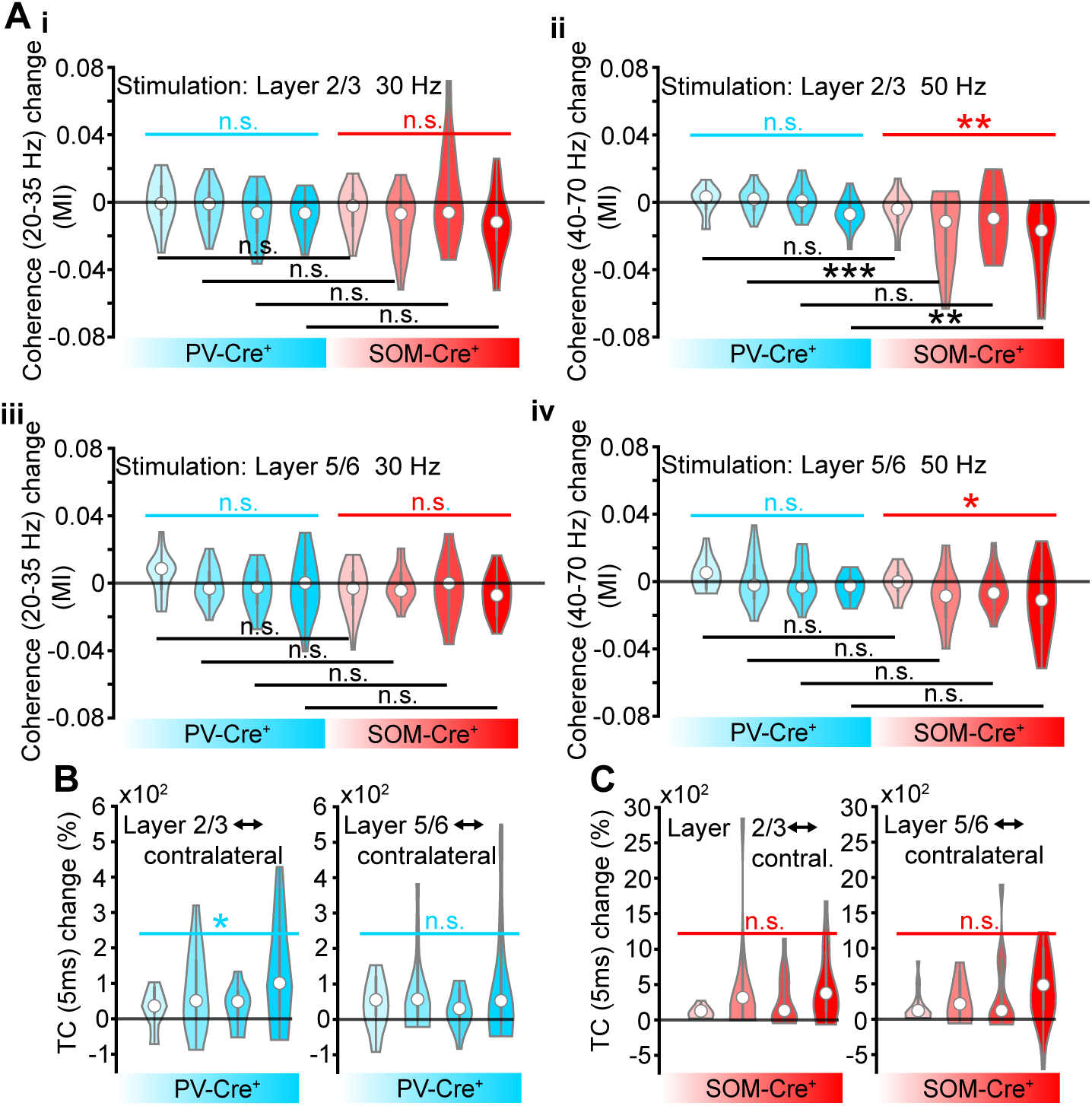
Role of PV+ and SOM+ interneurons for crosshemispheric communication. (**A**) Violinplots displaying the MI of imaginary coherence in the 20-35 Hz range between the baseline LFP and the LFP during optogenetic 30 Hz pulse stimulation in L2/3 (i) and L5/6 (iii) in the mPFC of PV-Cre^+^ and SOM-Cre^+^ and the MI of imaginary coherence in the 40-70 Hz range between the baseline LFP and the LFP during optogenetic 50 Hz pulse stimulation in L2/3 (ii) and L5/6 (iv) in the mPFC of PV-Cre^+^ (blue, n=83 recordings, 34 mice) and SOM-Cre^+^ (red, n=82 recordings, 33 mice) mice for each age group. (**B**) Violinplots displaying the percentage of TC change between baseline recording and optogenetic stimulation in the mPFC of PV-Cre^+^ mice for each age group. Left plot: TC is calculated between spike trains of ipsilateral L2/3 units and contralateral units (n=4940 unit pairs, 109 recordings, 53 mice); optogenetic pulse stimulation in L2/3. Right plot: TC is calculated between spike trains of ipsilateral L5/6 units and contralateral units (n=3687 unit pairs, 82 recordings, 32 mice); optogenetic pulse stimulation in L5/6. Time lag for TC calculation = 5ms. (**C**) Same as (C) for optogenetic stimulation of SOM-Cre^+^ mice (L2/3-contralateral: n=29104 unit pairs, 110 recordings, 53 mice; L5/6-contralateral: n=19524 unit pairs, 112 recordings, 49 mice). Violinplots in (A)-(C) represent the median with 25^th^ and 75^th^ percentile. In (A) - (C) colored asterisks indicate significant effect of age. * p < 0.05, ** p < 0.01, linear mixed-effect models. In (A) black asterisks indicate significant differences between stimulation of PV-Cre^+^ and SOM-Cre^+^ mice for all age groups. ** p < 0.01, *** p < 0.001, linear mixed-effect models. See Table S1 for detailed statistics.

Next, we calculated the TC with a 5 ms lag to elucidate the role of monosynaptic projections of PV+ and SOM+ INs for crosshemispheric communication. Stimulation of PV+ INs in L2/3 progressively increased the correlation between L2/3 units and units from the contralateral hemisphere along development. In contrast, no age-dependent effect on crosshemispheric correlation was detected after stimulation of PV+ INs in L5/6 or SOM+ INs in either layer (Fig. 4B+C). This suggests that with age PV+ INs in L2/3 increasingly contribute to crosshemispheric mPFC communication via direct projections.

## Discussion

Transient inhibitory circuits and resulting oscillatory patterns critically shape the maturation of various cortical regions. ^24,29,30,33^ Only if the assembly of neural networks from different cell types occurs in a temporally coordinated manner cognitive abilities can emerge at adult age. ^4,24,65^ Many cognitive functions, such as attention and working memory, rely on gamma oscillations and their crosshemispheric synchronization in the PFC ^3,6,48^, yet the developmental processes leading to the ability to generate and synchronize gamma oscillations across hemispheres are not fully uncovered.

We tracked bilateral prefrontal activity between pre-juvenile and adult age (P16-60) in mice to shine light on the physiological maturation and underlying cellular mechanisms of oscillatory synchrony within the local and crosshemispheric mPFC circuits. In addition to extracellular recordings, we optogenetically activated PV+ and SOM+ INs in a frequency and layer-specific manner. Considering, that oscillations in the mPFC reach frequencies above 50 Hz only at the end of the fourth postnatal week ^23^ and PV+ and SOM+ INs differ in their involvement in the generation of beta and gamma oscillations, ^44,66^, we carried out and compared optogenetic pulse stimulation with a frequency of 30 Hz and 50 Hz. We (i) observed increasing crosshemispheric gamma synchrony over age that is driven by L2/3. (ii) Following a similar timeline, the inhibitory function of PV+ INs in prefrontal networks emerges and PV+, but not SOM+, INs can operate at gamma frequencies. (iii) PV+, but not SOM+, INs show an age-dependent involvement in the transmission of gamma synchrony. Thus, the functional integration of PV+ INs in SOM+ dominated prefrontal circuits can be considered as a critical driver of the emergence of local prefrontal gamma power and crosshemispheric synchrony.

Local prefrontal activity decorrelates during the first two postnatal weeks. ^67^ Between the third postnatal week and adulthood, we did not observe major changes in the correlation of firing. Solely, FS-RS unit pairs increase in correlated firing, indicating the developmental embedding of FS PV+ INs as a mechanism underlying the increase in gamma synchrony. Crosshemispheric correlation of neural firing and the strength of the oscillatory drive (i.e., rhythmic neural input) originating in L2/3 decreases over age, while crosshemispheric gamma synchrony increases over age. This aligns with the general developmental phenomenon in which cortical activity becomes increasingly self-generated and less influenced by external inputs. ^34^ As a result, prefrontal crosshemispheric communication and information flow in the gamma range increase over age, whereas the underlying neural code sparsens, leading to less but optimized functional connections which more efficiently mediate gamma synchrony. Additionally, the increase in crosshemispheric gamma synchrony might also boost the crosshemispheric information exchange. ^47,68^ According to our findings, FS units in L2/3 play a central role for the synchronization of activity, since their firing is more correlated with contralateral units compared to RS and L5/6 units across age. Both prefrontal PV+ and SOM+ INs are targeted by long-range projections originating from the contralateral PFC. ^64^ However, only PV+ INs project to the contralateral PFC and have been shown to support crosshemispheric gamma synchronization. ^48^ Our data suggest that the age-dependent increase in gamma synchrony is mediated by an increase in long-range excitatory drive onto L2/3 PV+ INs as indicated by an increase in correlated firing between L2/3 PV+ INs and contralateral units.

In the mature PFC, PV+ and SOM+ INs show reciprocal inhibition and differ in their axonal targets (soma vs. dendrites), firing properties, and frequency specific generation of oscillation (gamma vs. beta) ^40,44^. We identified evidence for a shift from predominant inhibition of PV+ INs by SOM+ INs toward an increase in inhibition of SOM+ INs exerted by PV+ INs. This is in line with a previously reported increase in inhibitory inputs onto SOM+ INs between the first and fourth postnatal week. ^54^ The here identified functional integration of PV+ INs into SOM+ dominated circuits most likely leads to a switch in interneuronal dominance, causing faster inhibitory dynamics in prefrontal networks that are, in line with computational experiments ^20,35^, necessary for gamma generation. Correspondingly, with increasing prefrontal gamma synchrony along development, optogenetic 50 Hz stimulation of SOM+, but not PV+, INs in L2/3 causes a decrease in endogenous gamma power. Moreover, only PV+ INs show the ability to follow 50 Hz stimulation and exert efficient inhibition within gamma frequency ranges. This is in line with SOM+ INs mainly inhibiting other neurons’ dendrites, which is slower and temporally less precise compared to perisomatic inhibition exerted by PV+ INs. ^40,69^ Thus, 50 Hz stimulation of SOM+ INs probably resulted in out-of-frequency inhibition of PV+ INs and pyramidal neurons, ultimately disrupting endogenous gamma generation. Collectively our data suggest that from the third postnatal week on PV+ INs functionally operate in the gamma frequency range, whereas SOM+ INs fail to participate in gamma generation, confirming PV+ INs as driving force for the emergence of prefrontal gamma oscillations. However, the question remains why we did not observe an increase in oscillatory gamma activity upon 50 Hz activation of PV+ INs. This might be due to the absence of task-induced activity in this study since the generation and synchronization of gamma oscillations in the PFC is generally associated with cognitive behaviors. ^5,6,70^ PV+ INs most likely mainly act as organizers of excitatory activity to transmit gamma synchrony. Consequently, the lack of task-specific excitatory input could explain why 50 Hz stimulation of PV+ INs did not increase gamma synchrony.

Although it remains speculative why the prefrontal embedding of PV+ INs and the resulting emergence of gamma oscillations is protracted compared to other cortical areas, the upregulation of PV+ IN activity during defined periods is mandatory for mature cognitive abilities. ^17,65,71–73^ In sensory areas, early SOM+ IN mediated inhibition of PV+ INs and an increasing excitatory drive of pyramidal neurons onto PV+ INs regulate the activity-dependent maturation of PV+ INs. ^30,32,34,74,75^ Previous studies showed that disrupting early SOM+ IN mediated inhibition of PV+ INs or the activity of PV+ INs itself during adolescence causes impaired neural function in the mPFC and abnormal behavior later in life. ^17,30,48,65,71–73,75^ In contrast, studies focusing on the PFC revealed a developmental increase in PV expression, maturation of intrinsic firing properties of PV+ INs, and strengthening of connections between PV+ INs and L2/3 pyramidal neurons, reaching adult like levels during late adolescence. ^35,49–53,76,77^ SOM+ INs, on the other hand, do not undergo such age-related changes. ^23,54,78^ Studies in marmosets and humans rather showed a developmental decrease of SOM expression in the PFC. ^55,79^ In line with that, we found *in vivo* an increasing inhibitory effect of PV+, but not SOM+, INs in prefrontal networks, between the second postnatal week and adulthood. The onset of adolescence aligns with the increase in PV+ IN-mediated inhibition. This leads to the hypothesis that the rise of pubertal hormones might mediate the increase in PV+ IN activity. ^80^ However, our statistical analysis did not reveal any sex-related differences (Table S1) although puberty onset is delayed in males, indicating a rather minor impact of pubertal hormones on the exact timing of PV+ IN integration.

The importance of a precisely timed embedding of PV+ INs in prefrontal networks is further underpinned by the crucial role of PV+ INs in the pathophysiology of neurodevelopmental disorders, such as schizophrenia, where local prefrontal gamma power as well as crosshemispheric synchrony is disrupted. ^9,12,24,81–88^ By considering that the onset of disease symptoms during adolescence coincides with the emergence of mature gamma oscillations and the functional integration of PV+ INs, it is plausible that an altered integration of PV+ INs leads to the reported abnormal function of PV+ INs and the onset of cognitive disabilities. ^88^

Overall, our results provide experimental evidence for a hypothesized functional switch from SOM+ to PV+ INs ^27^, most likely controlling the emergence of mature gamma oscillations and crosshemispheric synchrony, mandatory for adult cognitive processing.

## Acknowledgments

We thank A. Dahlmann, A. Marquardt and P. Putthoff for excellent technical assistance.

## Author contributions

A.O., J.A.P., and I.L.H.-O. designed the experiments and interpreted the data. A.O. carried out experiments. A.O. performed data analysis. A.O., J.A.P., and I.L.H.-O. wrote the paper. All authors approved the submitted version.

## Supplemental information

Document S1. Figure legends S1-S2 and Table S1 (statistical results)

## STAR Methods

### Key resource table

| REAGENT or RESOURCE | SOURCE | IDENTIFIER |
| --- | --- | --- |
| **Antibodies** |  |  |
| Rabbit polyclonal primary antibody against GFP | Synaptic Systems | Cat# 132 002, RRID: AB_887725 |
| Alexa Fluor-488 goat anti-rabbit IgG secondary antibody | Thermo Fisher Scientific | Cat# A32731, RRID: AB_2633280 |
| DAPI BioChemica | PanReac AppliChem | #A1001 |
| **Bacterial and virus strains** |  |  |
| AAV9-Ef1alpha-DIO-hChR2(ET/TC)-eYFP | https://www.nature.com/articles/nmeth.1808 | Addgene viral prep # 35509-AAV9 |
| **Deposited data** |  |  |
| Extracellular recording data and sorted single unit data | This paper | Available upon request |
| Custom analysis code | This paper | / |
| **Experimental models: Organisms/strains** |  |  |
| Mouse: C57Bl/6J | Universitätsklinikum Hamburg-Eppendorf – Animal facility | Mouse: C57Bl/6J |
| Mouse: B6;129P2-*Pvalb*<sup>tm1(cre)Arbr</sup>/J (PV-Cre<sup>+</sup>) | Jackson Laboratory | RRID: IMSR_JAX:008069 |
| Mouse: *Sst*<sup>tm2.1(cre)Zjh</sup>/J (SOM-Cre<sup>+</sup>) | Jackson Laboratory | RRID: IMSR_JAX:013044 |
| **Software and algorithms** |  |  |
| Matlab 2017b, 2021b, | MathWorks | RRID:SCR_001622 |
| ImageJ 1.52b | ImageJ | RRID:SCR_003070 |
| R Statistical Software 4.2 | R Project for Statistical Computing | RRID:SCR_001905 |
| Cheetah 6 | Neuralynx | http://neuralynx.com/ |
| Anaconda 1.9.6 | Conda | RRID:SCR_018317 |
| phy | Cortex lab | https://github.com/cortex-lab/phy |
| Kilosort4 | https://www.nature.com/articles/s41592-024-02232-7 | https://github.com/MouseLand/Kilosort |
| Custom analysis code | This paper | / |
| **Other** |  |  |
| Arduino Uno SMD | Arduino | A000073 |
| Digital Lynx 4SX | Neuralynx | http://neuralynx.com/ |
| Diode laser (473 nm) | Omicron | LuxX® 473-100 |
| Recording optrode (4 shanks, 32 channels) | Neuronexus | A4x8-5mm-50-200-177-OA32LP |
| Recordings optrode (1 shank, 16 channels) | Neuronexus | A1x16-5mm-50-703-OA16LP |
| Recordings optrode (1 shank, 16 channels) | Neuronexus | A1x16-5mm-50-177-OA16LP |
| Microscope |  |  |
| Mobile HomeCage® | Neurotar | Mobile HomeCage Large® with tracking capability |
| Floating foam cage | Neurotar | Round cage; 325mm |
| Lightweight stainless-steel head plates | Neurotar | Model 3, round 6,2 mm opening |
| Headpost stand | Luigs & Neumann | 200-200 500 1100 |
| Head post - S (for mice) | Luigs & Neumann | 200-100 500 2203 |

### Resource availability

#### Lead contact

Ileana L. Hanganu-Opatz

#### Materials availability

The study did not generate new unique reagents.

#### Data and code availability

- LFP and SUA data reported in this paper will be shared by the lead contact upon request.
- All original or customized code will be deposited to an open-access repository and made publicly available. The DOI will be listed in the key resource table.
- Any additional information required to reanalyze the data reported in this paper is available from the lead contact upon request.

### Experimental model and study participant details

All experiments were performed in compliance with the German laws and the guidelines of the European Community for the use of animals in research and were approved by the local ethical committee (N19/121). Experiments were carried out on PV-Cre^+^, SOM-Cre^+^, and WT mice of both sexes. Cre^-^ littermates were used as (WT) control animals. Pregnant dams were housed individually. Weaned offspring was housed with at least two cage-mates on a 12 h light / 12 h dark cycle. The humidity and temperature were kept constant (40% relative humidity; 22°C). Food and water intake was not restricted. The day of vaginal plug detection was considered E0.5, the day of birth was considered P0. Experiments were performed on P1 to P60 mice.

## Method details

### Virus injections

At P1, 200 nl virus (titer, ≥ 1×10^13^ vg/mL) of Cre-dependent AAV9-Ef1alpha-DIO-hChR2(ET/TC)-eYFP (#35509, Addgene, MA, USA) ^89^ was injected into the mPFC (0.5 mm anterior to bregma, 0.1-0.5 mm lateral to the midline, 1.6 mm deep) of PV-/SOM-Cre^+^ and WT mice under isoflurane anesthesia (induction 5%, maintenance 2.5%). A NanoFil needle and syringe (NANOFIL, NF35BV, World Precision Instruments) connected to a micropump (Micro4, WPI) were used to inject the viral construct at a rate of 100 nl/min. During the injection, mice were placed on a heating blanket and left there until full recovery. The first recordings combined with optogenetic stimulation were performed not earlier than 15 days after injection, to enable reliable viral expression. (as described before ^25^)

### In vivo Electrophysiology

#### Chronic extracellular head-fixed recordings

Multisite extracellular head-fixed recordings were performed bilaterally in the mPFC of P16–17, P20-21, P30-33, and P50-60 mice. The adapter for head fixation was implanted at least 1 day before recordings. 30 minutes before surgery, buprenorphine (0.05 mg/kg body weight) was subcutaneously injected. Mice were anesthetized with isoflurane anesthesia (5% induction, 2.5% maintenance) and placed on a heating blanket to maintain the body temperature. Eyes were covered with eye ointment (Vidisic, Bausch + Lomb) to prevent drying. Depth of anesthesia and analgesia was evaluated with a toe pinch to test the paw withdrawal reflex. The skin of the head was disinfected with Betaisodona. After removing the skin, 0.5% bupivacaine/1% lidocaine was locally applied to wound edges. A metal head-post (Luigs and Neumann, Neurotar) was attached to the skull with dental cement and a craniotomy was performed above both mPFCs (1.75 mm anterior to bregma, 0.1–0.8 mm right/left to the midline) and protected by a customized synthetic window filled with Kwik-Cast sealant (World Precision Instruments). A silver wire was implanted between skull and brain tissue above the cerebellum and served as ground and reference. After surgery, mice were placed in a home cage on a heating blanket and returned to their mother/cage-mates after full recovery from anesthesia. Metacam (0.5 mg/ml, Boehringer-Ingelheim) mixed into soft food was provided for 2 days after surgery. For recordings, craniotomies were uncovered and multi-site optrodes (NeuroNexus) were stereotactically inserted into the right mPFC (one-shank, A4×8 recording sites, 50 μm spacing, 1.6 mm deep) and left mPFC (one-shank, A1×16 recording sites, 50 μm spacing, 1.8 mm deep). Recordings were acquired for 50-60 min. Extracellular signals were band-pass filtered (0.1–9000 Hz) and digitized (32 kHz) with a multichannel extracellular amplifier (Digital Lynx SX, Neuralynx). After recordings, the craniotomy was closed and mice were returned to their home cage as soon as possible. The same mice were recorded up to 5 times, but only within one age group. Optrode position was confirmed in brain slices postmortem. For LFP analysis of prefrontal activity in L2/3 and L5/6 of the right hemisphere as well as for the left hemisphere a recording channel within the PL was selected (as described before ^25^)

### Acute optogenetic stimulation

Optogenetic stimulation with short light pulses (3 ms) was performed using an Arduino uno controlled laser system (473 nm wavelength, Omicron). The laser was coupled with a 105 μm diameter light fiber (Thorlabs) to the optrodes. The integrated fiber of the optrodes ended 200 μm above the top recording site. 30 repetitions of stimulation with a pulse frequency of 30 and 50 Hz were conducted in L2/3 and L5/6 of the right mPFC. Laser power was adjusted to reliably induce neuronal firing and set between 2mW-6mW/mm at fiber tip. Responses to stimulations were averaged over 30 repetitions. PV-/SOM-Cre^+^ and WT mice underwent the same stimulation protocol.

### Histology

#### Perfusion and brain slicing

After the last recording, mice were anesthetized with 10% ketamine (aniMedica) / 2% xylazine (WDT) in 0.9% NaCl (10 μg/g body weight, intraperitoneal) and transcardially perfused with 4% paraformaldehyde (Histofix). Brains were taken out of the skull and postfixed in 4% paraformaldehyde for 24 h. Afterwards, brains were sectioned coronally with a vibratome at 100 μm for immunohistochemistry and reconstruction of optrode position.

### Immunohistochemistry

For immunohistochemistry, free-floating slices were washed 3 times for 15 min with PBS. Washed slices were permeabilized and blocked with PBS containing 0.75% Triton X-100 (Sigma-Aldrich, Darmstadt, Germany) and 5% normal goat serum (Cell Signaling Technology, Leiden, Netherlands) for 1 h. Slices were incubated over night with Rabbit polyclonal primary antibody against GFP (1:500, Cat# 132 002, RRID: AB_887725, Synaptic Systems) and afterwards washed for 3 times with PBS containing 0.2% Trition X-100. This was followed by 2 h of incubation with Alexa Fluor-488 goat anti-rabbit IgG secondary antibody (1:1000, Cat# A32731, RRID: AB_2633280, Thermo Fisher Scientific) and DAPI (5mg/ml, 1:2000, PanReac AppliChem) in PBS containing 0.5% Triton X-100 and 1% normal goat serum. After incubation, sections were 3 times washed for 15 min with PBS. Sections were transferred to glass slides and covered with Flouromount mounting medium (Thermo Fisher Scientific). (as described before ^25^)

## Quantification and statistical analysis

### Electrophysiological data analysis

#### LFP analysis

In vivo data were analyzed with custom-written algorithms in Matlab environment. 90 randomly selected 10 s-long signal segments were extracted from baseline recordings with a typical duration of 20 min and band-pass filtered (1–100 Hz) using a third-order Butterworth filter forward and backward to preserve phase information before down-sampling to analyze the LFP (as described before ^25^). To avoid that light artifacts during optogenetic stimulation might corrupt the LFP analysis, the 3 ms segments during the light pulse were replaced with a linear interpolation before band-pass filtering (1-100 Hz).

### Power spectral density

For power spectral density analysis 2 or 3 s-long windows of LFP signal (2 s: baseline, 3 s: pulse stimulation) were concatenated and the power was calculated using Welch’s method with non-overlapping windows.

### Imaginary coherence

The imaginary part of complex coherence, which is insensitive to volume conduction, was calculated by taking the absolute value of the imaginary component of the normalized cross-spectrum, ^58^ which was calculated between 1 s-long windows of LFP signal from two areas.

### Spectral density ratio (SDR)

To assess the directionality of causal influence between the LFP signals of two areas, we calculated the SDR ^60,61^ for both directions across the full frequency spectrum (1-100 Hz). Then we calculated the MI as (SDR 2 – SDR 1) / (SDR 2 + SDR 1) between SDR 1 (area 1 > area 2) and SDR 2 (area 1 < area 2), providing information about the direction and strength of causal influence.

### Generalized partial directed coherence (gPDC)

gPDC represents a frequency-domain representation of directionality based on the concept of Granger causality and was quantified on 1s-long LFP segments. The segments were denoised using the Matlab wavelet toolbox and further processed according to a previously described algorithm to quantify gPDC. ^59^ (as described before ^25^)

### Modulation index (MI)

For optogenetic stimulations, the modulation index was calculated as (power or coherence value during stimulation – power or coherence value during baseline recording) / (power or coherence value during stimulation + power or coherence value during baseline recording) for spectral densities of power and coherence of baseline recordings and 3 s-long segments during pulsed light stimulation.

### Single unit analysis

Spikes were detected with *kilosort* ^90^ and manually curated using *phy* (https://github.com/cortex-lab/phy). Data were imported and analyzed using custom-written routines in Matlab. The threshold for significantly modulated units during optogenetic pulse stimulation was set to an MI > 0 corresponding to activated and an MI < 0 to inhibited units and p < 0.05 (Wilcoxin signed-rank test). A time window of 5 ms pre and post stimulation onset was considered for positive modulation. For negative modulation the 5 ms pre stimulation were compared to the time window 6-10 ms post stimulation. To identify photo-tagged units the threshold was set to p < 0.001 and for MI to > 0.2. The duration of inhibition was calculated as the time units needed to reach their pre pulse firing rate (during the 5 ms before pulse onset) after pulse stimulation. In Fig. 2, for each recorded unit, if p-value of modulation after optogenetic stimulation in L2/3 was lower compared to stimulation in L5/6, modulation metrics were calculated for stimulation in L2/3 and vice versa.

### Classification of RS and FS units

RS and FS units were distinguished by setting a threshold based on spike halfwidth and trough to peak latency (halfwidth < 0.31 ms, trough to peak < 0.64 ms). It was previously shown that this threshold mainly identifies PV^+^ INs in L2/3 of the mPFC. ^24^ To assess how well both clusters separate, we calculated the average of the silhouette index ^91^ of all units using the built-in silhouette Matlab function. The silhouette index measures how similar each data point (i.e. unit) is to its own cluster compared to other clusters, providing a quantitative assessment of cluster separation.

### Tiling coefficient (TC)

TC, a firing rate-independent measure of correlation between two spike trains, was calculated according to a previously published algorithm. ^62^ For this, the proportion of spikes from spike train A that fall within ± Δt of a spike from spike train B within 5 or 10 ms is quantified, respectively. The resulting value is subtracted by the proportion of time that occurs within ± Δt of spikes from spike train B. This is then divided by 1 minus the product of these 2 values. The same is then applied after inverting spike train A and B, and the mean between the 2 values is kept which ranges between −1 and 1. (as described before ^25^)

### Model

Aiming to detect PV+ and SOM+ INs in our dataset of recorded single units, we trained two random forest classification models. ^92,93^ We extracted 8 spike-time based and 8 waveform based features, that were previously used to train a random forest classifier to detect PV+ INs by Sukman & Stark (2022) ^94^, for every recorded unit. Only the first part of each recording without optogenetic stimulation (baseline recording) was considered for calculation of spike-time based features. The features of units recorded in PV-Cre^+^ mice together with information about whether a unit was optogenetically phototagged (i.e. a PV+ IN) were used to train the PV+ classifier. Likewise, the SOM+ classifier was trained using features of units recorded in SOM-Cre^+^ mice. If a certain feature could not be calculated (e.g. because of too few spikes in the observed period), the value for this feature was set to the average value of all recorded units. As the portion of tagged units (4.3% in PV-Cre^+^ mice; 9.3% in SOM-Cre^+^ mice) was much smaller than the portion of not-tagged units, we assigned a class weight for each unit that was inversely proportional to the frequency of their class (tagged vs. not-tagged). This ensured that the classifier is not biased towards classifying the majority class. To train the classifiers, the datasets were split into a training and test dataset in an 80:20 ratio. A five-fold gridsearch was performed to find optimal hyperparameters for the classifier. We evaluated all possible combinations of the number of trees (set to 1, 10, 50, or 100) and the minimal number of samples at a leaf node (set to 1, 2, 4, 8, 16, or 32). The F1 score was calculated as a performance metric after each iteration and represents the harmonic mean between precision and recall of the model. ^95,96^ Afterwards the final classifiers were trained using the hyperparameters, which achieved the highest F1 score in the gridsearch.

To evaluate the overall performance of the classification procedure, we produced n=50 independent PV+ classifiers and n=50 SOM+ classifiers and calculated the corresponding F1 scores and AUCs (area under the receiver operating characteristics curve) ^97^. For comparison, we additionally trained each n=50 classifiers based on the datasets acquired in PV-Cre^+^ and in SOM-Cre^+^ mice with randomly mixed features and calculated their performance metrics.

### Statistical analysis

Statistical analyses were performed in R Statistical Software (Foundation for Statistical Computing) and Matlab (MathWorks). Considering that we used mice with different genotypes, sexes as well as acquired multiple recordings from one mouse, we used linear-mixed-effect models (*lmer* function of the *lme4* R package) for statistical comparison and Tukey multi comparison correction for post hoc analysis (*glht* function of the *multcomp* R package). Sex, mouse identity, mouse line and recording number where used as random parameters to calculate the linear-mixed effect models. The effect of sex and mouse line on the data distribution can be found in Table S1. Proportions of modulated units (Fig. 2C, S2A, S2C) were fitted using generalized linear-mixed effect models (*glmer* function of the *lme4* R package; Family = binomial) and confidence intervals were calculated using binomial models ^98^ to determine if SUA was modulated significantly (Fig. S2C). The Wilcoxin signed-rank test (built-in *signrank* function in Matlab) was used to test for significant differences of firing after 30 vs. 50 Hz pulse stimulation (Fig. 3B ii). Outlier removal was applied to data points if their distance from the 25th or 75th percentile exceeded 1.5 times the interquartile interval. See Table S1 for detailed statistical results.

---

**Figure S1. Related to Figure 1.**
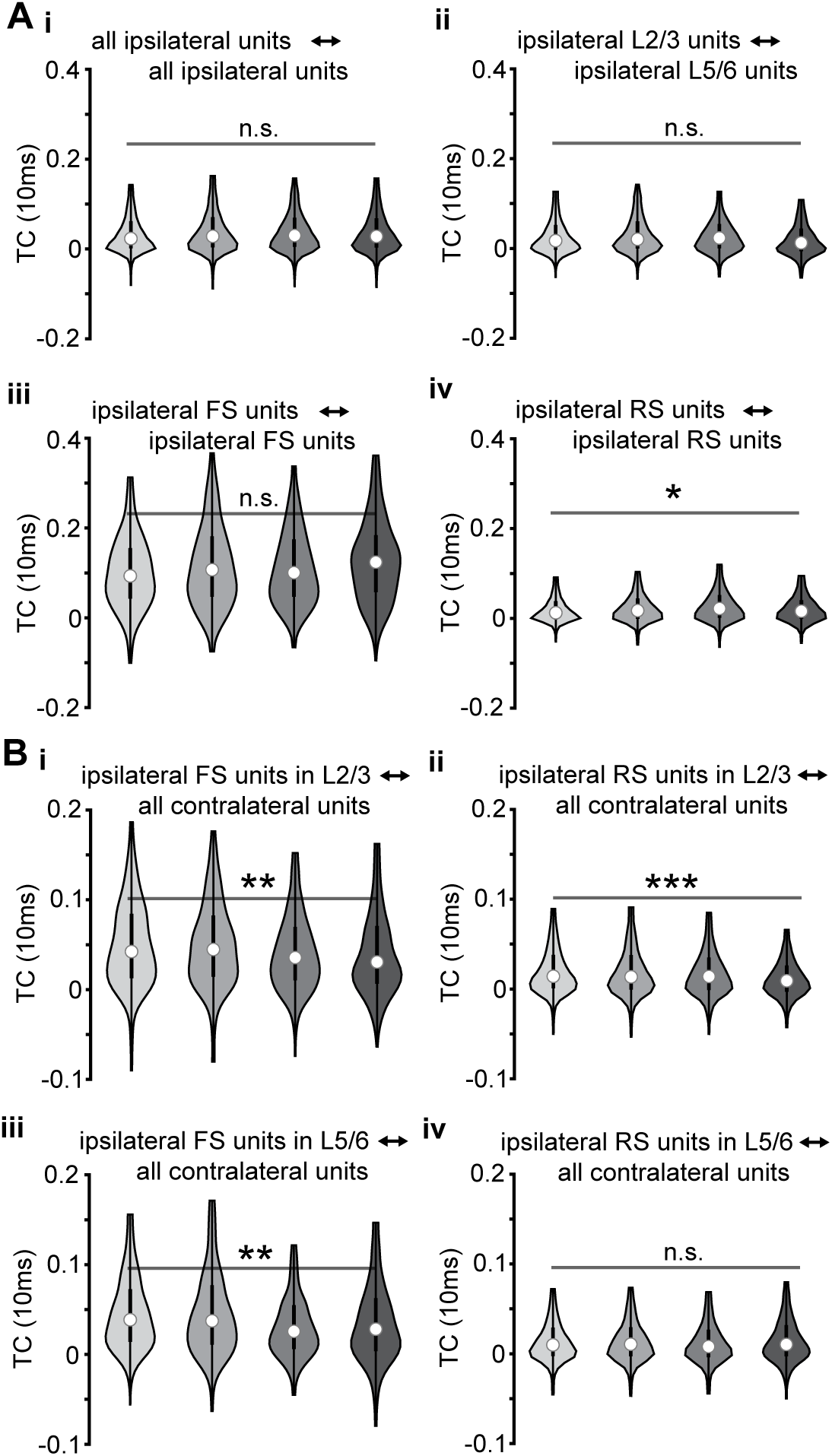
Local and crosshemispheric spike-time correlations

**Figure S2. Related to Figure 2.**
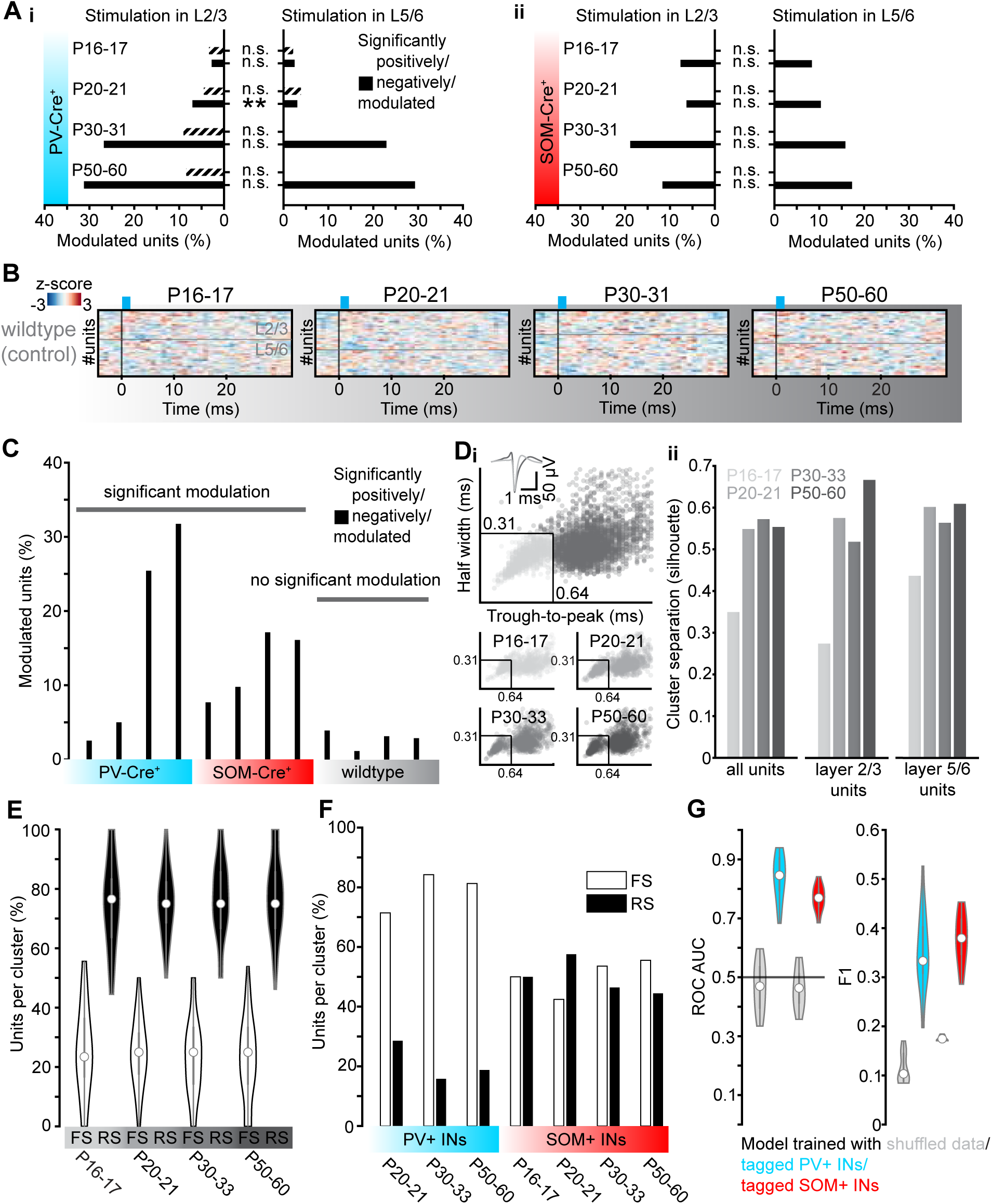
Stimulation of control animals, waveform based clustering of single units and model performance

