## Supplementary information for "Developmental embedding of parvalbumin interneurons drives local and crosshemispheric prefrontal gamma synchrony"

### Supplementary figure legends

**Figure S1. Related to Figure 1. Local and crosshemispheric spike-time correlations**

**(A)** Violinplots displaying the TC between simultaneously recorded spike trains of all ipsilateral units (i, n=28557 unit pairs, 236 recordings, 113 mice), ipsilateral L2/3 units and ipsilateral L5/6 units (ii, n=2130 unit pairs, 207 recordings, 111 mice), all ipsilateral FS units (iii, n=1695 unit pairs, 175 recordings, 101 mice), and all ipsilateral RS units (iv, n=16663 unit pairs, 232 recordings, 112 mice) for each age group. Time lag for TC calculation = 10ms.
**(B)** Violinplots displaying the TC between simultaneously recorded spike trains of ipsilateral FS units in L2/3 and contralateral units (i, n=3084 unit pairs, 150 recordings, 97 mice), ipsilateral RS units in L2/3 and contralateral units (ii, n=12313 unit pairs, 193 recordings, 104 mice), ipsilateral FS units in L5/6 and contralateral units (iii, n=4329 unit pairs, 155 recordings, 91 mice), and ipsilateral RS units in L5/6 and contralateral units (iv, n=12098 unit pairs, 196 recordings, 104 mice) for each age group. Time lag for TC calculation = 10ms.
Violinplots in (A + B) represent the median with $25^{th}$ and $75^{th}$ percentile. Asterisks indicate significant effect of age. * $p < 0.05$, ** $p < 0.01$, *** $p < 0.001$, linear mixed-effect models. See Table S1 for detailed statistics.

**Figure S2. Related to Figure 2. Stimulation of control animals, waveform based clustering of single units and model performance**

**(A)** Barplots displaying the absolute percentage of significantly modulated units after optogenetic 30 Hz pulse stimulation in L2/3 compared to L5/6 in PV-Cre$^+$ (i, n=110 recordings, 53 mice) and SOM-Cre$^+$ (ii, n=119 recordings, 52 mice) mice for each age group. Individual percentages were not calculated since binomial models were used for statistical analysis. Threshold for significant modulation $p < 0.05$. Asterisks indicate significant effect of layer of stimulation. ** $p < 0.01$
**(B)** Rasterplots showing the z-scored and period averaged spike trains of all recorded units in WT mice in each age group during optogenetic stimulation with blue (473 nm wavelength) 3 ms light pulses delivered in the ipsilateral mPFC with a frequency of 30 Hz. Units recorded in L2/3 are shown above and units recorded in L5/6 are shown below the grey horizontal lines. (P16-17 n=77 units, P20-21 n=78 units, P30-33 n=64 units, P50-60 n=69 units)
**(C)** Barplot displaying the absolute percentage of significantly modulated units after optogenetic 30 Hz pulse stimulation in L2/3 or L5/6 in PV-Cre$^+$ (n=1678 units, 53 mice), SOM-Cre$^+$ (n=1201 units, 50 mice), and WT (n=300 units, 12 mice) mice for each age group. Individual percentages were not calculated since binomial models were used for statistical analysis. Threshold for significant modulation $p < 0.05$.
**(D i)** Top, scatterplot showing half width and trough-to-peak latency of all ipsi- and contralaterally recorded units. Light grey units belong to FS cluster and dark grey units belong to RS cluster. Thresholds of the FS cluster (half width < 0.31 ms, trough-to-peak < 0.64 ms) are indicated by the box inset. Top left corner, mean waveforms of all FS and RS units. Bottom, same as top but showing all units of each age group. Plots were cut at half width = 0.6 and trough-to-peak = 1.5. (µV = microvolt)
**(D ii)** Barplots displaying the quality of the separation of the FS and RS cluster measured by the mean silhouette value of all single units for each age group. Left, including all ipsi-

and contralateral units. Middle, including all ipsilateral units in L2/3. Right, including all ipsilateral units in L5/6.

**(E)** Violinplots displaying the percentage of FS and RS units per animal for every age group (n=241 recordings, 115 mice).

**(F)** Barplots displaying the percentage of optogenetically tagged PV+ (n=110 recordings, 53 mice) and SOM+ (n=119 recordings, 50 mice) INs being FS and RS units for every age group.

**(G)** Left, violinplots displaying the AUC of the ROC curves of each 20 cross-validated random forest models based on a dataset including optogenetically tagged PV+ INs (blue), optogenetically tagged SOM+ INs (red), and correspondingly the dataset with optogenetically tagged PV+ INs but shuffled features, and the dataset with optogenetically tagged SOM+ INs but shuffled features (grey). Right, same as left, but quantifying the F1 score.

Violinplots in (E) and (G) represent the median with 25$^{th}$ and 75$^{th}$ percentile. See Table S1 for detailed statistics.

### Table S1

| Figure | Test | Factor | Groups | df | Test value | p-value | Male vs. female | Chi-Square | p-value | PV-Cre+ vs. SOM-Cre+ vs. WT | Chi-Square | p-value |
|---|---|---|---|---|---|---|---|---|---|---|---|---|
| **Figure 1** | | | | | | | | | | | | |
| Figure 1C i (bottom left) | Linear mixed effect model (LME) | Age | P16-P60 (110 mice, 236 recordings) | | | | Males: 125 recordings Females: 111 recordings | 0 | 1 | PV-Cre+: 103 recordings SOM-Cre+: 109 recordings wt: 24 recordings | 0 | 1 |
| | | | Imaginary coherence (20-35 Hz) | 3 | Chi-Square: 5.7434 | 0.1248 | | | | | | |
| | | | P16-17: (26 mice, 48 recordings) | | | | | | | | | |
| | | | P20-21: (29 mice, 52 recordings) | | | | | | | | | |
| | | | P30-33: (24 mice, 53 recordings) | | | | | | | | | |
| | | | P50-60: (31 mice, 83 recordings) | | | | | | | | | |
| | Tukey's post-hoc test of LME-model | Condition | P20-21 – P16-17 | | z-value: -0.920 | 1 | | | | | | |
| | | | P30-33 – P16-17 | | z-value: -2.239 | 0.151 | | | | | | |
| | | | P50-60 – P16-17 | | z-value: -1.698 | 0.448 | | | | | | |
| | | | P30-33 – P20-21 | | z-value: -1.378 | 0.673 | | | | | | |
| | | | P50-60 – P20-21 | | z-value: -0.760 | 1 | | | | | | |
| | | | P50-60 – P30-33 | | z-value: 0.713 | 1 | | | | | | |

| | | | | | | | | | | | | |
|---|---|---|---|---|---|---|---|---|---|---|---|---|
| Figure 1C i<br><br>(bottom right) | Linear mixed effect model (LME) | Age | P16-P60 (110 mice, 236 recordings) | | | | Males: 125 recordings Females: 111 recordings | 0.1804 | 0.6711 | PV-Cre+: 103 recordings SOM-Cre+: 109 recordings wt: 24 recordings | 0.3744 | 0.5406 |
| | | | Imaginary coherence (40-70 Hz) | 3 | Chi-Square: 1.4953 | 0.6834 | | | | | | |
| | | | P16-17: (26 mice, 48 recordings) | | | | | | | | | |
| | | | P20-21: (29 mice, 52 recordings) | | | | | | | | | |
| | | | P30-33: (24 mice, 53 recordings) | | | | | | | | | |
| | | | P50-60: (31 mice, 83 recordings) | | | | | | | | | |
| | Tukey's post-hoc test of LME-model | Condition | P20-21 – P16-17 | | z-value: 0.370 | 1 | | | | | | |
| | | | P30-33 – P16-17 | | z-value: 0.395 | 1 | | | | | | |
| | | | P50-60 – P16-17 | | z-value: 1.249 | 1 | | | | | | |
| | | | P30-33 – P20-21 | | z-value: 0.038 | 1 | | | | | | |
| | | | P50-60 – P20-21 | | z-value: 0.892 | 1 | | | | | | |
| | | | P50-60 – P30-33 | | z-value: 0.842 | 1 | | | | | | |
| Figure 1C ii<br><br>(bottom left) | Linear mixed effect model (LME) | Age | P16-P60 (110 mice, 236 recordings) | | | | Males: 125 recordings Females: 111 recordings | 0 | 1 | PV-Cre+: 103 recordings SOM-Cre+: 109 recordings wt: 24 recordings | 0 | 1 |
| | | | Imaginary coherence (20-35 Hz) | 3 | Chi-Square: 10.402 | 0.04132 | | | | | | |
| | | | P16-17: (26 mice, 48 recordings) | | | | | | | | | |
| | | | P20-21: (29 mice, 52 recordings) | | | | | | | | | |
| | | | P30-33: (24 mice, 53 recordings) | | | | | | | | | |
| | | | P50-60: (31 mice, 83 recordings) | | | | | | | | | |
| | Tukey's post-hoc test of LME-model | Condition | P20-21 – P16-17 | | z-value: 0.482 | 0.6297 | | | | | | |
| | | | P30-33 – P16-17 | | z-value: 2.711 | 0.0403 | | | | | | |
| | | | P50-60 – P16-17 | | z-value: 1.496 | 0.5385 | | | | | | |
| | | | P30-33 – P20-21 | | z-value: 2.297 | 0.1082 | | | | | | |
| | | | P50-60 – P20-21 | | z-value: 1.015 | 0.6205 | | | | | | |
| | | | P50-60 – P30-33 | | z-value: -1.450 | 0.5385 | | | | | | |
| Figure 1C ii | Linear mixed effect | Age | P16-P60 (110 mice, 236 recordings) | | | | | | | | | |

| | | | | | | | | | | | | |
|---|---|---|---|---|---|---|---|---|---|---|---|---|
| (bottom right) | model (LME) | | Imaginary coherence (40-70 Hz) | 3 | Chi-Square: 5.7434 | 0.01544 | | | | | | |
| | | | P16-17: (26 mice, 48 recordings) | | | | | | | | | |
| | | | P20-21: (29 mice, 52 recordings) | | | | | | | | | |
| | | | P30-33: (24 mice, 53 recordings) | | | | | | | | | |
| | | | P50-60: (31 mice, 83 recordings) | | | | | | | | | |
| | Tukey's post-hoc test of LME-model | Condition | P20-21 – P16-17 | | z-value: 0.187 | 0.8516 | | | | | | |
| | | | P30-33 – P16-17 | | z-value: 1.405 | 0.6403 | | | | | | |
| | | | P50-60 – P16-17 | | z-value: 2.783 | 0.0323 | | | | | | |
| | | | P30-33 – P20-21 | | z-value: 1.251 | 0.6403 | | | | | | |
| | | | P50-60 – P20-21 | | z-value: 2.656 | 0.0396 | | | | | | |
| | | | P50-60 – P30-33 | | z-value: 1.248 | 0.6403 | | | | | | |
| Figure 1C iii<br><br>(bottom left) | Linear mixed effect model (LME) | Age | P16-P60 (110 mice, 236 recordings) | | | | Males: 125 recordings Females: 111 recordings | 0 | 1 | PV-Cre+: 103 recordings SOM-Cre+: 109 recordings wt: 24 recordings | 0 | 1 |
| | | | Imaginary coherence (20-35 Hz) | 3 | Chi-Square: 3.0638 | 0.3819 | | | | | | |
| | | | P16-17: (26 mice, 48 recordings) | | | | | | | | | |
| | | | P20-21: (29 mice, 52 recordings) | | | | | | | | | |
| | | | P30-33: (24 mice, 53 recordings) | | | | | | | | | |
| | | | P50-60: (31 mice, 83 recordings) | | | | | | | | | |
| | Tukey's post-hoc test of LME-model | Condition | P20-21 – P16-17 | | z-value: -0.385 | 1 | | | | | | |
| | | | P30-33 – P16-17 | | z-value: 0.210 | 1 | | | | | | |
| | | | P50-60 – P16-17 | | z-value: -1.309 | 0.952 | | | | | | |
| | | | P30-33 – P20-21 | | z-value: 0.603 | 1 | | | | | | |
| | | | P50-60 – P20-21 | | z-value: -0.915 | 1 | | | | | | |
| | | | P50-60 – P30-33 | | z-value: -1.557 | 0.717 | | | | | | |
| Figure 1C iii<br><br>(bottom right) | Linear mixed effect model (LME) | Age | P16-P60 (110 mice, 236 recordings) | | | | Males: 125 recordings Females: 111 recordings | 0 | 1 | PV-Cre+: 103 recordings SOM-Cre+: 109 recordings wt: 24 recordings | 0 | 1 |
| | | | Imaginary coherence (40-70 Hz) | 3 | Chi-Square: 2.9379 | 0.4013 | | | | | | |
| | | | P16-17: (26 mice, 48 recordings) | | | | | | | | | |

| | | | | | | | | | | | | |
|---|---|---|---|---|---|---|---|---|---|---|---|---|
| | | | P20-21: (29 mice, 52 recordings) | | | | | | | | | |
| | | | P30-33: (24 mice, 53 recordings) | | | | | | | | | |
| | | | P50-60: (31 mice, 83 recordings) | | | | | | | | | |
| | Tukey's post-hoc test of LME-model | Condition | P20-21 – P16-17 | | z-value: 0.583 | 1 | | | | | | |
| | | | P30-33 – P16-17 | | z-value: 0.374 | 1 | | | | | | |
| | | | P50-60 – P16-17 | | z-value: 1.589 | 0.672 | | | | | | |
| | | | P30-33 – P20-21 | | z-value: -0.197 | 1 | | | | | | |
| | | | P50-60 – P20-21 | | z-value: 1.001 | 1 | | | | | | |
| | | | P50-60 – P30-33 | | z-value: 1.173 | 1 | | | | | | |
| Figure 1D i<br><br>(right) | Linear mixed effect model (LME) | Age | P16-P60 (111 mice, 234 recordings) | | | | Males: 123 recordings Females: 111 recordings | 0 | 1 | PV-Cre+: 101 recordings SOM-Cre+: 109 recordings wt: 24 recordings | 0.26 | 0.6101 |
| | | | gPDC (40-70 Hz) | 3 | Chi-Square: 10.747 | 0.01317 | | | | | | |
| | | | P16-17: (26 mice, 46 recordings) | | | | | | | | | |
| | | | P20-21: (29 mice, 52 recordings) | | | | | | | | | |
| | | | P30-33: (25 mice, 54 recordings) | | | | | | | | | |
| | | | P50-60: (31 mice, 82 recordings) | | | | | | | | | |
| | Tukey's post-hoc test of LME-model | Condition | P20-21 – P16-17 | | z-value: 0.583 | 0.38173 | | | | | | |
| | | | P30-33 – P16-17 | | z-value: 0.374 | 0.38173 | | | | | | |
| | | | P50-60 – P16-17 | | z-value: 1.589 | 0.00638 | | | | | | |
| | | | P30-33 – P20-21 | | z-value: -0.197 | 0.90854 | | | | | | |
| | | | P50-60 – P20-21 | | z-value: 1.001 | 0.33640 | | | | | | |
| | | | P50-60 – P30-33 | | z-value: 1.173 | 0.34331 | | | | | | |
| Figure 1D ii<br><br>(right) | Linear mixed effect model (LME) | Age | P16-P60 (111 mice, 234 recordings) | | | | Males: 123 recordings Females: 111 recordings | 0 | 1 | PV-Cre+: 101 recordings SOM-Cre+: 109 recordings wt: 24 recordings | 0 | 1 |
| | | | gPDC (40-70 Hz) | 3 | Chi-Square: 19.579 | 0.0002075 | | | | | | |
| | | | P16-17: (26 mice, 46 recordings) | | | | | | | | | |
| | | | P20-21: (29 mice, 52 recordings) | | | | | | | | | |
| | | | P30-33: (25 mice, 54 recordings) | | | | | | | | | |
| | | | P50-60: (31 mice, 82 recordings) | | | | | | | | | |

| | | | | | | | | | | | | |
|---|---|---|---|---|---|---|---|---|---|---|---|---|
| | Tukey's post-hoc test of LME-model | Condition | P20-21 – P16-17 | | z-value: 0.583 | 0.944392 | | | | | | |
| | | | P30-33 – P16-17 | | z-value: 0.374 | 0.007168 | | | | | | |
| | | | P50-60 – P16-17 | | z-value: 1.589 | 0.000338 | | | | | | |
| | | | P30-33 – P20-21 | | z-value: -0.197 | 0.037661 | | | | | | |
| | | | P50-60 – P20-21 | | z-value: 1.001 | 0.003380 | | | | | | |
| | | | P50-60 – P30-33 | | z-value: 1.173 | 0.944392 | | | | | | |
| Figure 1D iii<br><br>(right) | Linear mixed effect model (LME) | Age | P16-P60 (111 mice, 234 recordings) | | | | Males: 123 recordings Females: 111 recordings | 0 | 1 | PV-Cre+: 101 recordings SOM-Cre+: 109 recordings wt: 24 recordings | 0.0009 | 0.9764 |
| | | | gPDC (40-70 Hz) | 3 | Chi-Square: 16.45 | 0.0009167 | | | | | | |
| | | | P16-17: (26 mice, 46 recordings) | | | | | | | | | |
| | | | P20-21: (29 mice, 52 recordings) | | | | | | | | | |
| | | | P30-33: (25 mice, 54 recordings) | | | | | | | | | |
| | | | P50-60: (31 mice, 82 recordings) | | | | | | | | | |
| | Tukey's post-hoc test of LME-model | Condition | P20-21 – P16-17 | | z-value: 0.181 | 0.85608 | | | | | | |
| | | | P30-33 – P16-17 | | z-value: 1.762 | 0.30951 | | | | | | |
| | | | P50-60 – P16-17 | | z-value: 3.607 | 0.00186 | | | | | | |
| | | | P30-33 – P20-21 | | z-value: 1.634 | 0.30951 | | | | | | |
| | | | P50-60 – P20-21 | | z-value: 3.536 | 0.00203 | | | | | | |
| | | | P50-60 – P30-33 | | z-value: 1.766 | 0.30951 | | | | | | |
| Figure 1E | Linear mixed effect model (LME) | Age | P16-P60 (110 mice, 233 recordings) | | | | Males: 124 recordings Females: 109 recordings | 0 | 1 | PV-Cre+: 103 recordings SOM-Cre+: 106 recordings wt: 24 recordings | 0 | 1 |
| | | | SDR MI | 3 | Chi-Square: 29.121 | 0.000002112 | | | | | | |
| | | | P16-17: (26 mice, 48 recordings) | | | | | | | | | |
| | | | P20-21: (29 mice, 52 recordings) | | | | | | | | | |
| | | | P30-33: (24 mice, 53 recordings) | | | | | | | | | |
| | | | P50-60: (31 mice, 80 recordings) | | | | | | | | | |
| | Tukey's post-hoc test of | Condition | P20-21 – P16-17 | | z-value: 3.598 | 0.001281 | | | | | | |
| | | | P30-33 – P16-17 | | z-value: -0.642 | 0.808976 | | | | | | |

| | | | | | | | | | | | | |
|---|---|---|---|---|---|---|---|---|---|---|---|---|
| | LME-model | | P50-60 – P16-17 | | z-value: -1.511 | 0.392606 | | | | | | |
| | | | P30-33 – P20-21 | | z-value: -4.207 | 0.000129 | | | | | | |
| | | | P50-60 – P20-21 | | z-value: -5.354 | 0.000000517 | | | | | | |
| | | | P50-60 – P30-33 | | z-value: -0.834 | 0.808976 | | | | | | |
| Figure 1F | Linear mixed effect model (LME) | Age | P16-P60 (110 mice, 233 recordings) | | | | Males: 124 recordings Females: 109 recordings | 0 | 1 | PV-Cre+: 103 recordings SOM-Cre+: 106 recordings wt: 24 recordings | 0 | 1 |
| | | | SDR MI | 3 | Chi-Square: 29.784 | 0.000001532 | | | | | | |
| | | | P16-17: (26 mice, 48 recordings) | | | | | | | | | |
| | | | P20-21: (29 mice, 52 recordings) | | | | | | | | | |
| | | | P30-33: (24 mice, 53 recordings) | | | | | | | | | |
| | | | P50-60: (31 mice, 80 recordings) | | | | | | | | | |
| | Tukey's post-hoc test of LME-model | Condition | P20-21 – P16-17 | | z-value: 3.763 | 0.000672 | | | | | | |
| | | | P30-33 – P16-17 | | z-value: -0.735 | 0.924495 | | | | | | |
| | | | P50-60 – P16-17 | | z-value: -1.295 | 0.586106 | | | | | | |
| | | | P30-33 – P20-21 | | z-value: -4.460 | 4.09e-05 | | | | | | |
| | | | P50-60 – P20-21 | | z-value: -5.301 | 6.92e-07 | | | | | | |
| | | | P50-60 – P30-33 | | z-value: -0.514 | 0.924495 | | | | | | |
| Figure 1G | Linear mixed effect model (LME) | Age | P16-P60 (108 mice, 5216 unit pairs) | | | | Males: 3131 unit pairs Females: 2085 unit pairs | 0 | 1 | PV-Cre+: 2943 unit pairs SOM-Cre+: 1861 unit pairs wt: 412 unit pairs | 0 | 1 |
| | | | Tiling coefficient (10ms) | 3 | Chi-Square: 7.9765 | 0.0465 | | | | | | |
| | | | P16-17: (25 mice, 1101 unit pairs) | | | | | | | | | |
| | | | P20-21: (29 mice, 1558 unit pairs) | | | | | | | | | |
| | | | P30-33: (25 mice, 972 unit pairs) | | | | | | | | | |
| | | | P50-60: (29 mice, 1585 unit pairs) | | | | | | | | | |
| | Tukey's post-hoc test of LME-model | Condition | P20-21 – P16-17 | | z-value: 1.362 | 0.5195 | | | | | | |
| | | | P30-33 – P16-17 | | z-value: 1.646 | 0.4985 | | | | | | |
| | | | P50-60 – P16-17 | | z-value: 2.789 | 0.0318 | | | | | | |
| | | | P30-33 – P20-21 | | z-value: 0.385 | 0.7000 | | | | | | |

| | | | | | | | | | | | | |
|---|---|---|---|---|---|---|---|---|---|---|---|---|
| | | | P50-60 – P20-21 | | z-value: 1.526 | 0.5077 | | | | | | |
| | | | P50-60 – P30-33 | | z-value: 1.056 | 0.5815 | | | | | | |
| Figure 1H FS | Linear mixed effect model (LME) | Age | P16-P60 (102 mice, 7420 unit pairs) | | | | Males: 4376 unit pairs Females: 3044 unit pairs | 0.4861 | 0.4857 | PV-Cre+: 4457 unit pairs SOM-Cre+: 2503 unit pairs wt: 460 unit pairs | 0 | 1 |
| | | | Tiling coefficient (10ms) | 3 | Chi-Square: 20.837 | 0.0001138 | | | | | | |
| Figure 1H RS | Linear mixed effect model (LME) | Age | P16-P60 (105 mice, 24489 unit pairs) | | | | Males: 14698 unit pairs Females: 9791 unit pairs | 0 | 1 | PV-Cre+: 13879 unit pairs SOM-Cre+: 8437 unit pairs wt: 2173 unit pairs | 0 | 1 |
| | | | Tiling coefficient (10ms) | 3 | Chi-Square: 7.6463 | 0.05392 | | | | | | |
| Figure 1H | Tukey's post-hoc test of LME-model | Condition | FS - RS (P16-17, 27 mice, 6725 unit pairs) | | z-value: 39.699 | 0.0000000 | | | | | | |
| | | | FS - RS (P20-21, 26 mice, 6319 unit pairs) | | z-value: 35.513 | 0.0000000 | | | | | | |
| | | | FS - RS (P30-33, 23 mice, 7983 unit pairs) | | z-value: 26.275 | 0.0000000 | | | | | | |
| | | | FS - RS (P50-60, 30 mice, 10882 unit pairs) | | z-value: 35.392 | 0.0000000 | | | | | | |
| Figure 1I L2/3 | Linear mixed effect model (LME) | Age | P16-P60 (102 mice, 15337 unit pairs) | | | | Males: 9243 unit pairs Females: 6094 unit pairs | 0 | 1 | PV-Cre+: 8647 unit pairs SOM-Cre+: 5518 unit pairs wt: 1172 unit pairs | 0 | 1 |
| | | | Tiling coefficient (10ms) | 3 | Chi-Square: 25.016 | 0.00001532 | | | | | | |
| Figure 1I L5/6 | Linear mixed effect model (LME) | Age | P16-P60 (105 mice, 16510 unit pairs) | | | | Males: 9839 unit pairs Females: 6671 unit pairs | 0 | 1 | PV-Cre+: 9636 unit pairs SOM-Cre+: 5414 unit pairs wt: 1460 unit pairs | 0 | 1 |
| | | | Tiling coefficient (10ms) | 3 | Chi-Square: 10.547 | 0.01444 | | | | | | |
| Figure 1I | Tukey's post-hoc test of LME-model | Condition | L23 - L5/6 (P16-17, 27 mice, 6713 unit pairs) | | z-value: 3.164 | 0.029522 | | | | | | |
| | | | L23 - L5/6 (P20-21, 26 mice, 6355 unit pairs) | | z-value: 10.545 | 0.0000000 | | | | | | |
| | | | L23 - L5/6 (P30-33, 23 mice, 7983 unit pairs) | | z-value: 8.297 | 0.0000000 | | | | | | |

| | | | | | | | | | | | | |
|---|---|---|---|---|---|---|---|---|---|---|---|---|
| | | | L23 - L5/6 (P50-60, 30 mice, 10796 unit pairs) | | z-value: -1.840 | 0.723995 | | | | | | |
| **Figure S1** | | | | | | | | | | | | |
| Figure S1A i | Linear mixed effect model (LME) | Age | P16-P60 (113 mice, 28557 unit pairs) | | | | Males: 16392 unit pairs Females: 12165 unit pairs | 0 | 1 | PV-Cre+: 15948 unit pairs SOM-Cre+: 10269 unit pairs wt: 2340 unit pairs | 0 | 1 |
| | | | Tiling coefficient (10ms) | 3 | Chi-Square: 2.2554 | 0.5211 | | | | | | |
| | | | P16-17: (27 mice, 6745 unit pairs) | | | | | | | | | |
| | | | P20-21: (29 mice, 8614 unit pairs) | | | | | | | | | |
| | | | P30-33: (26 mice, 5650 unit pairs) | | | | | | | | | |
| | | | P50-60: (31 mice, 7548 unit pairs) | | | | | | | | | |
| | Tukey's post-hoc test of LME-model | Condition | P20-21 – P16-17 | | z-value: 1.100 | 1 | | | | | | |
| | | | P30-33 – P16-17 | | z-value: 1.283 | 1 | | | | | | |
| | | | P50-60 – P16-17 | | z-value: 0.370 | 1 | | | | | | |
| | | | P30-33 – P20-21 | | z-value: 0.248 | 1 | | | | | | |
| | | | P50-60 – P20-21 | | z-value: -0.739 | 1 | | | | | | |
| | | | P50-60 – P30-33 | | z-value: -0.947 | 1 | | | | | | |
| Figure S1A ii | Linear mixed effect model (LME) | Age | P16-P60 (111 mice, 2130 unit pairs) | | | | Males: 1041 unit pairs Females: 1089 unit pairs | 0 | 1 | PV-Cre+: 1152 unit pairs SOM-Cre+: 856 unit pairs wt: 122 unit pairs | 0.1974 | 0.6568 |
| | | | Tiling coefficient (10ms) | 3 | Chi-Square: 4.3193 | 0.229 | | | | | | |
| | | | P16-17: (25 mice, 546 unit pairs) | | | | | | | | | |
| | | | P20-21: (29 mice, 479 unit pairs) | | | | | | | | | |
| | | | P30-33: (26 mice, 541 unit pairs) | | | | | | | | | |
| | | | P50-60: (29 mice, 564 unit pairs) | | | | | | | | | |
| | Tukey's post-hoc test of LME-model | Condition | P20-21 – P16-17 | | z-value: 0.490 | 1 | | | | | | |
| | | | P30-33 – P16-17 | | z-value: -0.301 | 1 | | | | | | |
| | | | P50-60 – P16-17 | | z-value: -1.652 | 0.493 | | | | | | |
| | | | P30-33 – P20-21 | | z-value: -0.775 | 1 | | | | | | |
| | | | P50-60 – P20-21 | | z-value: -2.176 | 0.177 | | | | | | |
| | | | P50-60 – P30-33 | | z-value: -1.297 | 0.779 | | | | | | |

| | | | | | | | | | | | | |
|---|---|---|---|---|---|---|---|---|---|---|---|---|
| Figure S1A iii | Linear mixed effect model (LME) | Age | P16-P60 (101 mice, 1695 unit pairs) | | | | Males: 895 unit pairs Females: 800 unit pairs | 0.0089 | 0.9247 | PV-Cre+: 1002 unit pairs SOM-Cre+: 591 unit pairs wt: 102 unit pairs | 0 | 1 |
| | | | Tiling coefficient (10ms) | 3 | Chi-Square: 4.9979 | 0.1719 | | | | | | |
| | | | P16-17: (24 mice, 467 unit pairs) | | | | | | | | | |
| | | | P20-21: (28 mice, 493 unit pairs) | | | | | | | | | |
| | | | P30-33: (23 mice, 238 unit pairs) | | | | | | | | | |
| | | | P50-60: (26 mice, 497 unit pairs) | | | | | | | | | |
| | Tukey's post-hoc test of LME-model | Condition | P20-21 – P16-17 | | z-value: 1.381 | 0.670 | | | | | | |
| | | | P30-33 – P16-17 | | z-value: 1.536 | 0.622 | | | | | | |
| | | | P50-60 – P16-17 | | z-value: 2.242 | 0.150 | | | | | | |
| | | | P30-33 – P20-21 | | z-value: 0.305 | 1 | | | | | | |
| | | | P50-60 – P20-21 | | z-value: 0.972 | 0.993 | | | | | | |
| | | | P50-60 – P30-33 | | z-value: 0.591 | 1 | | | | | | |
| Figure S1A iv | Linear mixed effect model (LME) | Age | P16-P60 (112 mice, 16663 unit pairs) | | | | Males: 9622 unit pairs Females: 7041 unit pairs | 0 | 1 | PV-Cre+: 9249 unit pairs SOM-Cre+: 5909 unit pairs wt: 1505 unit pairs | 0 | 1 |
| | | | Tiling coefficient (10ms) | 3 | Chi-Square: 10.232 | 0.01669 | | | | | | |
| | | | P16-17: (26 mice, 3802 unit pairs) | | | | | | | | | |
| | | | P20-21: (29 mice, 4986 unit pairs) | | | | | | | | | |
| | | | P30-33: (26 mice, 3603 unit pairs) | | | | | | | | | |
| | | | P50-60: (31 mice, 4272 unit pairs) | | | | | | | | | |
| | Tukey's post-hoc test of LME-model | Condition | P20-21 – P16-17 | | z-value: 0.817 | 1 | | | | | | |
| | | | P30-33 – P16-17 | | z-value: 2.970 | 0.0179 | | | | | | |
| | | | P50-60 – P16-17 | | z-value: 0.429 | 1 | | | | | | |
| | | | P30-33 – P20-21 | | z-value: 2.262 | 0.0947 | | | | | | |
| | | | P50-60 – P20-21 | | z-value: -0.391 | 1 | | | | | | |
| | | | P50-60 – P30-33 | | z-value: -2.616 | 0.0445 | | | | | | |
| Figure S1B i | Linear mixed effect | Age | P16-P60 (97 mice, 3084 unit pairs) | | | | Males: 1870 unit pairs | 0 | 1 | PV-Cre+: 1789 unit pairs SOM-Cre+: 1092 unit | 0 | 1 |

| | | | | | | | | | | | | |
|---|---|---|---|---|---|---|---|---|---|---|---|---|
| | model (LME) | | Tiling coefficient (10ms) | 3 | Chi-Square: 11.544 | 0.00912 | Females: 1214 unit pairs | | | pairs wt: 203 unit pairs | | |
| | | | P16-17: (24 mice, 581 unit pairs) | | | | | | | | | |
| | | | P20-21: (5 mice, 641 unit pairs) | | | | | | | | | |
| | | | P30-33: (21 mice, 803 unit pairs) | | | | | | | | | |
| | | | P50-60: (27 mice, 1059 unit pairs) | | | | | | | | | |
| | Tukey's post-hoc test of LME-model | Condition | P20-21 – P16-17 | | z-value: -0.393 | 1 | | | | | | |
| | | | P30-33 – P16-17 | | z-value: -2.392 | 0.0758 | | | | | | |
| | | | P50-60 – P16-17 | | z-value: -2.766 | 0.0341 | | | | | | |
| | | | P30-33 – P20-21 | | z-value: -2.064 | 0.1171 | | | | | | |
| | | | P50-60 – P20-21 | | z-value: -2.428 | 0.0758 | | | | | | |
| | | | P50-60 – P30-33 | | z-value: -0.190 | 1 | | | | | | |
| Figure S1B ii | Linear mixed effect model (LME) | Age | P16-P60 (104 mice, 12313 unit pairs) | | | | Males: 7390 unit pairs Females: 4923 unit pairs | 0 | 1 | PV-Cre+: 6865 unit pairs SOM-Cre+: 4477 unit pairs wt: 971 unit pairs | 0 | 1 |
| | | | Tiling coefficient (10ms) | 3 | Chi-Square: 16.596 | 0.0008557 | | | | | | |
| | | | P16-17: (26 mice, 2619 unit pairs) | | | | | | | | | |
| | | | P20-21: (23 mice, 2602 unit pairs) | | | | | | | | | |
| | | | P30-33: (23 mice, 3370 unit pairs) | | | | | | | | | |
| | | | P50-60: (29 mice, 3722 unit pairs) | | | | | | | | | |
| | Tukey's post-hoc test of LME-model | Condition | P20-21 – P16-17 | | z-value: 0.074 | 0.94093 | | | | | | |
| | | | P30-33 – P16-17 | | z-value: -1.550 | 0.31952 | | | | | | |
| | | | P50-60 – P16-17 | | z-value: -3.550 | 0.00192 | | | | | | |
| | | | P30-33 – P20-21 | | z-value: -1.614 | 0.31952 | | | | | | |
| | | | P50-60 – P20-21 | | z-value: -3.607 | 0.00186 | | | | | | |
| | | | P50-60 – P30-33 | | z-value: -1.854 | 0.25492 | | | | | | |
| Figure S1B iii | Linear mixed effect model (LME) | Age | P16-P60 (91 mice, 4329 unit pairs) | | | | Males: 2509 unit pairs Females: 1820 unit pairs | 0.3267 | 0.5676 | PV-Cre+: 2655 unit pairs SOM-Cre+: 1415 unit pairs wt: 259 unit pairs | 0 | 1 |
| | | | Tiling coefficient (10ms) | 3 | Chi-Square: 12.281 | 0.006481 | | | | | | |
| | | | P16-17: (24 mice, 1180 unit pairs) | | | | | | | | | |

| | | | | | | | | | | | | |
|---|---|---|---|---|---|---|---|---|---|---|---|---|
| | | | P20-21: (22 mice, 779 unit pairs) | | | | | | | | | |
| | | | P30-33: (18 mice, 924 unit pairs) | | | | | | | | | |
| | | | P50-60: (27 mice, 1446 unit pairs) | | | | | | | | | |
| | Tukey's post-hoc test of LME-model | Condition | P20-21 – P16-17 | | z-value: 0.916 | 0.7189 | | | | | | |
| | | | P30-33 – P16-17 | | z-value: -2.293 | 0.0874 | | | | | | |
| | | | P50-60 – P16-17 | | z-value: -1.951 | 0.1532 | | | | | | |
| | | | P30-33 – P20-21 | | z-value: -3.070 | 0.0128 | | | | | | |
| | | | P50-60 – P20-21 | | z-value: -2.803 | 0.0253 | | | | | | |
| | | | P50-60 – P30-33 | | z-value: 0.475 | 0.7189 | | | | | | |
| Figure S1B iv | Linear mixed effect model (LME) | Age | P16-P60 (104 mice, 12098 unit pairs) | | | | Males: 7346 unit pairs Females: 4852 unit pairs | 0.1234 | 0.7254 | PV-Cre+: 7035 unit pairs SOM-Cre+: 3961 unit pairs wt: 1202 unit pairs | 0 | 1 |
| | | | Tiling coefficient (10ms) | 3 | Chi-Square: 0.1028 | 0.9915 | | | | | | |
| | | | P16-17: (25 mice, 1101 unit pairs) | | | | | | | | | |
| | | | P20-21: (29 mice, 1558 unit pairs) | | | | | | | | | |
| | | | P30-33: (25 mice, 972 unit pairs) | | | | | | | | | |
| | | | P50-60: (29 mice, 1585 unit pairs) | | | | | | | | | |
| | Tukey's post-hoc test of LME-model | Condition | P20-21 – P16-17 | | z-value: 0.089 | 1 | | | | | | |
| | | | P30-33 – P16-17 | | z-value: -0.235 | 1 | | | | | | |
| | | | P50-60 – P16-17 | | z-value: -0.127 | 1 | | | | | | |
| | | | P30-33 – P20-21 | | z-value: -0.317 | 1 | | | | | | |
| | | | P50-60 – P20-21 | | z-value: -0.216 | 1 | | | | | | |
| | | | P50-60 – P30-33 | | z-value: 0.119 | 1 | | | | | | |
| **Figure 2** | | | | | | | | | | | | |
| Figure 2C PV-Cre+ | P16-17, 12 mice, 317 units, 17 positively modulated, 17 negatively modulated | | | | | | | | | | | |
| | P20-21, 13 mice, 481 units, 33 positively modulated, 48 negatively modulated | | | | | | | | | | | |
| | P30-33, 15 mice, 354 units, 42 positively modulated, 123 negatively modulated | | | | | | | | | | | |
| | P50-60, 13 mice, 526 units, 61 positively modulated, 213 negatively modulated | | | | | | | | | | | |
| PV-Cre+ positive modulation | Generalized linear mixed effect model | Age | Positively modulated units | 3 | Chi-Square: 12.922 | 0.004809 | Males: 877 units Females: | 0 | 1 | | | |

| | | | | | | | | | |
|---|---|---|---|---|---|---|---|---|---|
| | (GLME, binomial) | | | | | | 801 unit pairs | | |
| | Tukey's post-hoc test of GLME-model | Condition | P20-21 – P16-17 | | z-value: 0.649 | 0.9981 | | | |
| | | | P30-33 – P16-17 | | z-value: 2.491 | 0.1973 | | | |
| | | | P50-60 – P16-17 | | z-value: 2.619 | 0.1469 | | | |
| | | | P30-33 – P20-21 | | z-value: 2.121 | 0.3978 | | | |
| | | | P50-60 – P20-21 | | z-value: 2.269 | 0.3081 | | | |
| | | | P50-60 – P30-33 | | z-value: 0.053 | 1 | | | |
| PV-Cre+ negative modulation | Generalized linear mixed effect model (GLME, binomial) | Age | Positively modulated units | 3 | Chi-Square: 28.044 | 3.555e-06 | Males: 877 units Females: 801 units | 0 | 1 |
| | Tukey's post-hoc test of GLME-model | Condition | P20-21 – P16-17 | | z-value: 1.382 | 0.86439 | | | |
| | | | P30-33 – P16-17 | | z-value: 4.665 | < 0.001 | | | |
| | | | P50-60 – P16-17 | | z-value: 5.010 | < 0.001 | | | |
| | | | P30-33 – P20-21 | | z-value: 3.712 | 0.00504 | | | |
| | | | P50-60 – P20-21 | | z-value: 4.116 | < 0.001 | | | |
| | | | P50-60 – P30-33 | | z-value: 0.442 | 0.99985 | | | |
| Figure 2C SOM-Cre+ | P16-17, 13 mice, 325 units, 55 positively modulated, 42 negatively modulated | | | | | | | | |
| | P20-21, 13 mice, 317 units, 53 positively modulated, 41 negatively modulated | | | | | | | | |
| | P30-33, 9 mice, 292 units, 57 positively modulated, 69 negatively modulated | | | | | | | | |
| | P50-60, 15 mice, 267 units, 39 positively modulated, 63 negatively modulated | | | | | | | | |
| SOM-Cre+ positive modulation | Generalized linear mixed effect model (GLME, binomial) | Age | Positively modulated units | 3 | Chi-Square: 1.9477 | 0.5833 | Males: 666 units Females: 535 units | 0 | 0.9999 |
| | Tukey's post-hoc test of GLME-model | Condition | P20-21 – P16-17 | | z-value: 0.265 | 1 | | | |
| | | | P30-33 – P16-17 | | z-value: 1.041 | 0.9677 | | | |
| | | | P50-60 – P16-17 | | z-value: -0.435 | 0.9999 | | | |

| | | | | | | | | | |
|---|---|---|---|---|---|---|---|---|---|
| | | | P30-33 – P20-21 | | z-value: 0.791 | 0.9935 | | | |
| | | | P50-60 – P20-21 | | z-value: -0.691 | 0.9972 | | | |
| | | | P50-60 – P30-33 | | z-value: -1.425 | 0.8442 | | | |
| SOM-Cre+ negative modulation | Generalized linear mixed effect model (GLME, binomial) | Age | Positively modulated units | 3 | Chi-Square: 5.3953 | 0.145 | Males: 666 units Females: 535 units | 0 | 1 |
| | Tukey's post-hoc test of GLME-model | Condition | P20-21 – P16-17 | | z-value: 0.118 | 1 | | | |
| | | | P30-33 – P16-17 | | z-value: 1.652 | 0.71628 | | | |
| | | | P50-60 – P16-17 | | z-value: 1.425 | 0.84476 | | | |
| | | | P30-33 – P20-21 | | z-value: 1.543 | 0.78251 | | | |
| | | | P50-60 – P20-21 | | z-value: 1.306 | 0.89588 | | | |
| | | | P50-60 – P30-33 | | z-value: -0.346 | 0.99997 | | | |
| Figure 2D<br><br>PV-Cre+ | Linear mixed effect model (LME) | Age | P16-P60 (41 mice, 289 units) | | | | Males: 132 units Females: 157 units | 0 | 1 |
| | | | Inhibition strength (MI) | 3 | Chi-Square: 12.151 | 0.006883 | | | |
| | | | P16-17: (6 mice, 8 units) | | | | | | |
| | | | P20-21: (10 mice, 24 units) | | | | | | |
| | | | P30-33: (13 mice, 90 units) | | | | | | |
| | | | P50-60: (12 mice, 167 units) | | | | | | |
| | Tukey's post-hoc test of LME-model | Condition | P20-21 – P16-17 | | z-value: -0.265 | 1 | | | |
| | | | P30-33 – P16-17 | | z-value: -0.873 | 1 | | | |
| | | | P50-60 – P16-17 | | z-value: -2.572 | 0.0486 | | | |
| | | | P30-33 – P20-21 | | z-value: -0.770 | 1 | | | |
| | | | P50-60 – P20-21 | | z-value: -3.004 | 0.0160 | | | |
| | | | P50-60 – P30-33 | | z-value: -2.586 | 0.0486 | | | |
| Figure 2D<br><br>SOM-Cre+ | Linear mixed effect | Age | P16-P60 (33mice, 149 unit pairs) | | | | Males: 70 units | 0 | 1 |

| | | | | | | | | | |
|---|---|---|---|---|---|---|---|---|---|
| | model (LME) | | Inhibition strength (MI) | 3 | Chi-Square: 4.3959 | 0.2218 | Females: 79 units | | |
| | | | P16-17: (8 mice, 25 units) | | | | | | |
| | | | P20-21: 9 mice, 31 units) | | | | | | |
| | | | P30-33: (7 mice, 50 units) | | | | | | |
| | | | P50-60: (9 mice, 43 units) | | | | | | |
| | Tukey's post-hoc test of LME-model | Condition | P20-21 – P16-17 | | z-value: -0.962 | 1 | | | |
| | | | P30-33 – P16-17 | | z-value: 0.504 | 1 | | | |
| | | | P50-60 – P16-17 | | z-value: -1.186 | 0.943 | | | |
| | | | P30-33 – P20-21 | | z-value: 1.505 | 0.661 | | | |
| | | | P50-60 – P20-21 | | z-value: -0.222 | 0.1 | | | |
| | | | P50-60 – P30-33 | | z-value: -1.765 | 0.466 | | | |
| Figure 2E<br>PV-Cre+ | Linear mixed effect model (LME) | Age | P16-P60 (41 mice, 289 units) | | | | Males: 132 units Females: 157 units | 0 | 1 |
| | | | Inhibition duration (ms) | 3 | Chi-Square: 15.382 | 0.001518 | | | |
| | | | P16-17: (6 mice, 8 units) | | | | | | |
| | | | P20-21: (10 mice, 24 units) | | | | | | |
| | | | P30-33: (13 mice, 90 units) | | | | | | |
| | | | P50-60: (12 mice, 167 units) | | | | | | |
| | Tukey's post-hoc test of LME-model | Condition | P20-21 – P16-17 | | z-value: -0.488 | 1 | | | |
| | | | P30-33 – P16-17 | | z-value: 0.448 | 1 | | | |
| | | | P50-60 – P16-17 | | z-value: 2.098 | 0.143487 | | | |
| | | | P30-33 – P20-21 | | z-value: 1.389 | 0.494525 | | | |
| | | | P50-60 – P20-21 | | z-value: 3.786 | 0.000918 | | | |
| | | | P50-60 – P30-33 | | z-value: 3.000 | 0.013491 | | | |
| Figure 2E<br>SOM-Cre+ | Linear mixed effect model (LME) | Age | P16-P60 (33mice, 149 unit pairs) | | | | Males: 70 units Females: 79 units | 0 | 1 |
| | | | Inhibition duration (ms) | 3 | Chi-Square: 10.939 | 0.01206 | | | |
| | | | P16-17: (8 mice, 25 units) | | | | | | |

| | | | | | | |
|---|---|---|---|---|---|---|
| | | | P20-21: 9 mice, 31 units) | | | |
| | | | P30-33: (7 mice, 50 units) | | | |
| | | | P50-60: (9 mice, 43 units) | | | |
| | Tukey's post-hoc test of LME-model | Condition | P20-21 – P16-17 | | z-value: 0.407 | 1 |
| | | | P30-33 – P16-17 | | z-value: 0.730 | 1 |
| | | | P50-60 – P16-17 | | z-value: 2.901 | 0.0223 |
| | | | P30-33 – P20-21 | | z-value: 0.304 | 1 |
| | | | P50-60 – P20-21 | | z-value: 2.633 | 0.0404 |
| | | | P50-60 – P30-33 | | z-value: 2.649 | 0.0404 |
| Figure 2F PV-Cre+ | P16-17, 12 mice, 21 recordings | | | | | |
| | P20-21, 13 mice, 25 recordings | | | | | |
| | P30-33, 15 mice, 28 recordings | | | | | |
| | P50-60, 13 mice, 33 recordings | | | | | |
| PV-Cre+ Inhibited units (%) | Tukey's post-hoc test of LME-model | Condition | P16-17: FS - RS | | z-value: 0.007 | 1 |
| | | | P20-21: FS - RS | | z-value: -0.615 | 1 |
| | | | P30-33: FS - RS | | z-value: -2.134 | 1 |
| | | | P50-60: FS - RS | | z-value: -0.132 | 1 |
| Figure 2F SOM-Cre+ | P16-17, 13 mice, 24 recordings | | | | | |
| | P20-21, 13 mice, 24 recordings | | | | | |
| | P30-33, 9 mice, 23 recordings | | | | | |
| | P50-60, 15 mice, 35 recordings | | | | | |
| SOM-Cre+ Inhibited units (%) | Tukey's post-hoc test of LME-model | Condition | P16-17: FS - RS | | z-value: 1.014 | 1 |
| | | | P20-21: FS - RS | | z-value: 2.625 | 0.615584 |
| | | | P30-33: FS - RS | | z-value: 4.543 | 0.000666 |
| | | | P50-60: FS - RS | | z-value: 2.707 | 0.550324 |
| Figure 2G ii (top) | % of inhibited SOM+ units | P16-17: 4.92% | | | | |
| | | P20-21: 12.64% | | | | |
| | | P30-33: 33.82% | | | | |
| | | P50-60: 53.01% | | | | |
| Figure 2G ii (bottom) | | P16-17: 57.14% | | | | |
| | | P20-21: 48.00% | | | | |

| | | | | | | |
|---|---|---|---|---|---|---|
| | % of inhibited PV+ units | P30-33: 57.14% | | | | |
| | | P50-60: 25.09% | | | | |
| **Figure S2** | | | | | | |
| Figure S2A i (PV-Cre+) | P16-17, 12 mice, 317 units, 10 positively modulated L2/3, 9 negatively modulated L2/3, 7 positively modulated L5/6, 8 negatively modulated L5/6 | | | | | |
| | P20-21, 13 mice, 481 units, 22 positively modulated L2/3, 33 negatively modulated L2/3, 19 positively modulated L5/6, 15 negatively modulated L5/6 | | | | | |
| | P30-33, 15 mice, 354 units, 33 positively modulated L2/3, 96 negatively modulated L2/3, 23 positively modulated L5/6, 83 negatively modulated L5/6 | | | | | |
| | P50-60, 13 mice, 526 units, 45 positively modulated L2/3, 167 negatively modulated L2/3, 36 positively modulated L5/6, 152 negatively modulated L5/6 | | | | | |
| PV-Cre+ positive modulation | Tukey's post-hoc test of GLME-model | Layer of stimulation (L2/3 vs. L5/6) | P16-17 | | z-value: 0.734 | 0.4631 |
| | | | P20-21 | | z-value: 0.482 | 0.6296 |
| | | | P30-33 | | z-value: 1.386 | 0.1659 |
| | | | P50-60 | | z-value: 1.071 | 0.2840 |
| PV-Cre+ negative modulation | Tukey's post-hoc test of GLME-model | Layer of stimulation (L2/3 vs. L5/6) | P16-17 | | z-value: 0.246 | 0.8059 |
| | | | P20-21 | | z-value: 2.694 | 0.0071 |
| | | | P30-33 | | z-value: 1.205 | 0.2282 |
| | | | P50-60 | | z-value: 1.142 | 0.2534 |
| Figure S2A ii (SOM-Cre+) | P16-17, 12 mice, 325 units, 43 positively modulated L2/3, 25 negatively modulated L2/3, 33 positively modulated L5/6, 28 negatively modulated L5/6 | | | | | |
| | P20-21, 13 mice, 317 units, 35 positively modulated L2/3, 21 negatively modulated L2/3, 40 positively modulated L5/6, 31 negatively modulated L5/6 | | | | | |
| | P30-33, 15 mice, 292 units, 35 positively modulated L2/3, 56 negatively modulated L2/3, 37 positively modulated L5/6, 49 negatively modulated L5/6 | | | | | |
| | P50-60, 13 mice, 267 units, 33 positively modulated L2/3, 35 negatively modulated L2/3, 29 positively modulated L5/6, 47 negatively modulated L5/6 | | | | | |
| SOM-Cre+ positive modulation | Tukey's post-hoc test of GLME-model | Layer of stimulation (L2/3 vs. L5/6) | P16-17 | | z-value: 1.270 | 0.2042 |
| | | | P20-21 | | z-value: -0.636 | 0.5249 |
| | | | P30-33 | | z-value: -0.263 | 0.7927 |

| | | | | | | |
|---|---|---|---|---|---|---|
| | | | P50-60 | | z-value: 0.553 | 0.5805 |
| SOM-Cre+ negative modulation | Tukey's post-hoc test of GLME-model | Layer of stimulation (L2/3 vs. L5/6) | P16-17 | | z-value: -0.442 | 0.6585 |
| | | | P20-21 | | z-value: -1.530 | 0.1260 |
| | | | P30-33 | | z-value: 0.851 | 0.3949 |
| | | | P50-60 | | z-value: -1.516 | 0.1294 |
| Figure S2C | CI: 95% confidence interval for a binomial model (if the CI includes values lower than 0.05, modulation of SUA was not significant) | PV-Cre+ P16-17, 12 mice, 317 units, proportion of modulated units: 0.07886435 CI [0.05398665 0.1138266] | | | | |
| | | PV-Cre+ P20-21, 13 mice, 481 units, proportion of modulated units: 0.1185031 CI [0.0925985 0.150453] | | | | |
| | | PV-Cre+ P30-33, 15 mice, 354 units, proportion of modulated units: 0.3728814 CI [0.3241245 0.4243675] | | | | |
| | | PV-Cre+ P50-60, 13 mice, 526 units, proportion of modulated units: 0.4334601 CI [0.3917444 0.4761406] | | | | |
| | | SOM-Cre+ P16-17, 13 mice, 325 units, proportion of modulated units: 0.2461538 CI [0.2024663 0.2957721] | | | | |
| | | SOM-Cre+ P20-21, 13 mice, 317 units, proportion of modulated units: 0.2649842 CI [0.2194257 0.3161704] | | | | |
| | | SOM-Cre+ P30-33, 9 mice, 292 units, proportion of modulated units: 0.3664384 CI [0.3132401 0.4231052] | | | | |
| | | SOM-Cre+ P50-60, 15 mice, 267 units, proportion of modulated units: 0.3071161 CI [0.2548459 0.3648578] | | | | |
| | | wildtype P16-17, 2 mice, 77 units, proportion of modulated units: 0.06493506 CI [0.02805308 0.1431643] | | | | |
| | | wildtype P20-21, 4 mice, 89 units, proportion of modulated units: 0.05617978 CI [0.02423288 0.1248542] | | | | |
| | | wildtype P30-33, 2 mice, 64 units, proportion of modulated units: 0.0625 CI [0.0245712 0.1499749] | | | | |
| | | wildtype P50-60, 4 mice, 70 units, proportion of modulated units: 0.02857143 CI [0.007870612 0.09832256] | | | | |
| Figure S2D ii (left) | Silhouette index all units | P16-17: 0.35 | | | | |
| | | P20-21: 0.55 | | | | |
| | | P30-33: 0.57 | | | | |
| | | P50-60: 0.55 | | | | |
| Figure S2D ii (middle) | Silhouette index L2/3 units | P16-17: 0.27 | | | | |
| | | P20-21: 0.58 | | | | |
| | | P30-33: 0.52 | | | | |
| | | P50-60: 0.67 | | | | |
| | | P16-17: 0.44 | | | | |

| | | | | | | |
|---|---|---|---|---|---|---|
| Figure S2D ii (right) | Silhouette index L5/6 units | P20-21: 0.60 | | | | |
| | | P30-33: 0.56 | | | | |
| | | P50-60: 0.61 | | | | |
| Figure S2F | % in FS cluster | P16-17 SOM+ INs: 50% | | | | |
| | | P20-21 SOM+ INs: 42.42% PV+ INs: 71.43% | | | | |
| | | P30-33 SOM+ INs: 53.57% PV+ INs: 84.21% | | | | |
| | | P50-60 SOM+ INs: 55.56% PV+ INs: 81.25% | | | | |
| **Figure 3** | | | | | | |
| Figure 3B i PV-Cre+ | P16-17, 12 mice, 21 recordings | | | | | |
| | P20-21, 13 mice, 25 recordings | | | | | |
| | P30-33, 15 mice, 28 recordings | | | | | |
| | P50-60, 13 mice, 32 recordings | | | | | |
| PV-Cre+ Activated units (%) | Tukey's post-hoc test of LME-model | Condition | P16-17: 30 Hz - 50 Hz | | z-value: 0.360 | 1 |
| | | | P20-21: 30 Hz - 50 Hz | | z-value: -0.211 | 1 |
| | | | P30-33: 30 Hz - 50 Hz | | z-value: 0.177 | 1 |
| | | | P50-60: 30 Hz - 50 Hz | | z-value: -0.529 | 1 |
| Figure 3B i SOM-Cre+ | P16-17, 13 mice, 22 recordings | | | | | |
| | P20-21, 13 mice, 24 recordings | | | | | |
| | P30-33, 9 mice, 24 recordings | | | | | |
| | P50-60, 15 mice, 36 recordings | | | | | |
| SOM-Cre+ Activated units (%) | Tukey's post-hoc test of LME-model | Condition | P16-17: 30 Hz - 50 Hz | | z-value: -1.996 | 1 |
| | | | P20-21: 30 Hz - 50 Hz | | z-value: -0.539 | 1 |
| | | | P30-33: 30 Hz - 50 Hz | | z-value: 0.196 | 1 |
| | | | P50-60: 30 Hz - 50 Hz | | z-value: -0.987 | 1 |
| Figure 3B ii PV-Cre+ | P16-17, 12 mice, 21 recordings | | | | | |
| | P20-21, 13 mice, 25 recordings | | | | | |
| | P30-33, 15 mice, 28 recordings | | | | | |
| | P50-60, 13 mice, 32 recordings | | | | | |
| PV-Cre+ Activated units MI 30 vs. 50 Hz | Wilcoxin signed-rank test | Condition | P16-17: 30 Hz - 50 Hz | | p=0.0651 | |
| | | | P20-21: 30 Hz - 50 Hz | | p=0.5797 | |
| | | | P30-33: 30 Hz - 50 Hz | | p=0.8802 | |
| | | | P50-60: 30 Hz - 50 Hz | | p=0.0851 | |

| | | | | | | | | | |
|---|---|---|---|---|---|---|---|---|---|
| Figure 3B ii SOM-Cre+ | P16-17, 13 mice, 22 recordings | | | | | | | | |
| | P20-21, 13 mice, 24 recordings | | | | | | | | |
| | P30-33, 9 mice, 24 recordings | | | | | | | | |
| | P50-60, 15 mice, 36 recordings | | | | | | | | |
| SOM-Cre+ Activated units MI 30 vs. 50 Hz | Wilcoxin signed-rank test | Condition | P16-17: 30 Hz - 50 Hz | | p<0.0001 | | | | |
| | | | P20-21: 30 Hz - 50 Hz | | p=0.006 | | | | |
| | | | P30-33: 30 Hz - 50 Hz | | p<0.0001 | | | | |
| | | | P50-60: 30 Hz - 50 Hz | | p=0.0002 | | | | |
| Figure 3B iii PV-Cre+ | P16-17, 12 mice, 21 recordings | | | | | | | | |
| | P20-21, 12 mice, 24 recordings | | | | | | | | |
| | P30-33, 15 mice, 28 recordings | | | | | | | | |
| | P50-60, 13 mice, 33 recordings | | | | | | | | |
| PV-Cre+ Inhibited units (%) | Tukey's post-hoc test of LME-model | Condition | P16-17: 30 Hz - 50 Hz | | z-value: 0.283 | 1 | | | |
| | | | P20-21: 30 Hz - 50 Hz | | z-value: 0.090 | 1 | | | |
| | | | P30-33: 30 Hz - 50 Hz | | z-value: 0.452 | 1 | | | |
| | | | P50-60: 30 Hz - 50 Hz | | z-value: 2.311 | 1 | | | |
| Figure 3B iii SOM-Cre+ | P16-17, 13 mice, 24 recordings | | | | | | | | |
| | P20-21, 12 mice, 22 recordings | | | | | | | | |
| | P30-33, 9 mice, 24 recordings | | | | | | | | |
| | P50-60, 15 mice, 36 recordings | | | | | | | | |
| SOM-Cre+ Inhibited units (%) | Tukey's post-hoc test of LME-model | Condition | P16-17: 30 Hz - 50 Hz | | z-value: 1.690 | 1 | | | |
| | | | P20-21: 30 Hz - 50 Hz | | z-value: 0.949 | 1 | | | |
| | | | P30-33: 30 Hz - 50 Hz | | z-value: 5.873 | 4.85e-07 | | | |
| | | | P50-60: 30 Hz - 50 Hz | | z-value: 5.243 | 1.71e-05 | | | |
| Figure 3C i PV-Cre+ | Linear mixed effect model (LME) | Age | P16-P60 (35 mice, 102 recordings) | | | | Males: 49 recordings Females: 53 recordings | 0 | 1 |
| | | | Power 20-35 Hz (MI) | 3 | Chi-Square: 2.0613 | 0.5598 | | | |
| Figure 3C i SOM-Cre+ | Linear mixed effect | Age | P16-P60 (36 mice, 105 recordings) | | | | Males: 59 recordings Females: | 0 | 1 |

| | | | | | | | | | |
|---|---|---|---|---|---|---|---|---|---|
| | model (LME) | | Power 20-35 Hz (MI) | 3 | Chi-Square: 10.129 | 0.0175 | 46 recordings | | |
| Figure 3C i | Tukey's post-hoc test of LME-model | Condition | P16-17: PV-Cre+ - SOM-Cre+ | | z-value: 2.669 | 0.129387 | | | |
| | | | P20-21: PV-Cre+ - SOM-Cre+ | | z-value: 3.624 | 0.006670 | | | |
| | | | P30-33: PV-Cre+ - SOM-Cre+ | | z-value: 0.777 | 1 | | | |
| | | | P50-60: PV-Cre+ - SOM-Cre+ | | z-value: 2.539 | 0.177916 | | | |
| Figure 3C ii<br>PV-Cre+ | Linear mixed effect model (LME) | Age | P16-P60 (35 mice, 102 recordings) | | | | Males: 48 recordings Females: 54 recordings | 0 | 1 |
| | | | Power 40-70 Hz (MI) | 3 | Chi-Square: 19.031 | 0.0002694 | | | |
| Figure 3C ii<br>SOM-Cre+ | Linear mixed effect model (LME) | Age | P16-P60 (36 mice, 105 recordings) | | | | Males: 58 recordings Females: 47 recordings | 0 | 1 |
| | | | Power 40-70 Hz (MI) | 3 | Chi-Square: 5.4587 | 0.1411 | | | |
| Figure 3C ii | Tukey's post-hoc test of LME-model | Condition | P16-17: PV-Cre+ - SOM-Cre+ | | z-value: 2.091 | 0.328601 | | | |
| | | | P20-21: PV-Cre+ - SOM-Cre+ | | z-value: 7.700 | 3.52e-13 | | | |
| | | | P30-33: PV-Cre+ - SOM-Cre+ | | z-value: 5.881 | 8.57e-08 | | | |
| | | | P50-60: PV-Cre+ - SOM-Cre+ | | z-value: 5.345 | 1.81e-06 | | | |
| Figure 3C iii<br>PV-Cre+ | Linear mixed effect model (LME) | Age | P16-P60 (36 mice, 103 recordings) | | | | Males: 48 recordings Females: 55 recordings | 0 | 1 |
| | | | Power 20-35 Hz (MI) | 3 | Chi-Square: 3.1592 | 0.3677 | | | |
| Figure 3C iii<br>SOM-Cre+ | Linear mixed effect model (LME) | Age | P16-P60 (35 mice, 104 recordings) | | | | Males: 57 recordings Females: 47 recordings | 0 | 1 |
| | | | Power 20-35 Hz (MI) | 3 | Chi-Square: 3.3411 | 0.342 | | | |
| Figure 3C iii | Tukey's post-hoc test of | Condition | P16-17: PV-Cre+ - SOM-Cre+ | | z-value: -0.846 | 1 | | | |

| | | | | | | | | | |
|---|---|---|---|---|---|---|---|---|---|
| | LME-model | | P20-21: PV-Cre+ - SOM-Cre+ | | z-value: 2.029 | 1 | | | |
| | | | P30-33: PV-Cre+ - SOM-Cre+ | | z-value: 1.251 | 1 | | | |
| | | | P50-60: PV-Cre+ - SOM-Cre+ | | z-value: -0.640 | 1 | | | |
| Figure 3C iv<br><br>PV-Cre+ | Linear mixed effect model (LME) | Age | P16-P60 (36 mice, 100 recordings) | | | | Males: 47 recordings Females: 53 recordings | 0 | 1 |
| | | | Power 40-70 Hz (MI) | 3 | Chi-Square: 14.123 | 0.002742 | | | |
| Figure 3C iv<br><br>SOM-Cre+ | Linear mixed effect model (LME) | Age | P16-P60 (36 mice, 105 recordings) | | | | Males: 59 recordings Females: 46 recordings | 0.2138 | 0.6438 |
| | | | Power 40-70 Hz (MI) | 3 | Chi-Square: 4.7764 | 0.1889 | | | |
| Figure 3C iv | Tukey's post-hoc test of LME-model | Condition | P16-17: PV-Cre+ - SOM-Cre+ | | z-value: 1.285 | 1 | | | |
| | | | P20-21: PV-Cre+ - SOM-Cre+ | | z-value: 3.772 | 0.00356 | | | |
| | | | P30-33: PV-Cre+ - SOM-Cre+ | | z-value: 2.794 | 0.09901 | | | |
| | | | P50-60: PV-Cre+ - SOM-Cre+ | | z-value: 4.020 | 0.00152 | | | |
| **Figure 4** | | | | | | | | | |
| Figure 4A i<br><br>PV-Cre+ | Linear mixed effect model (LME) | Age | P16-P60 (34 mice, 83 recordings) | | | | Males: 42 recordings Females: 41 recordings | 0 | 1 |
| | | | Imaginary coherence (20-35 Hz) | 3 | Chi-Square: 5.2451 | 0.1547 | | | |
| Figure 4A i<br><br>SOM-Cre+ | Linear mixed effect model (LME) | Age | P16-P60 (33 mice, 82 recordings) | | | | Males: 47 recordings Females: 35 recordings | 0 | 1 |
| | | | Imaginary coherence (20-35 Hz) | 3 | Chi-Square: 3.7147 | 0.294 | | | |
| Figure 4A i | Tukey's post-hoc test of LME-model | Condition | P16-17: PV-Cre+ - SOM-Cre+ | | z-value: 0.802 | 1 | | | |
| | | | P20-21: PV-Cre+ - SOM-Cre+ | | z-value: 2.364 | 0.452 | | | |

| | | | | | | | | | |
|---|---|---|---|---|---|---|---|---|---|
| | | | P30-33: PV-Cre+ - SOM-Cre+ | | z-value: -1.375 | 1 | | | |
| | | | P50-60: PV-Cre+ - SOM-Cre+ | | z-value: 1.283 | 1 | | | |
| Figure 4A ii<br><br>PV-Cre+ | Linear mixed effect model (LME) | Age | P16-P60 (32 mice, 80 recordings) | | | | Males: 42 recordings Females: 38 recordings | 0 | 1 |
| | | | Imaginary coherence (40-70 Hz) | 3 | Chi-Square: 3.2876 | 0.3494 | | | |
| Figure 4A ii<br><br>SOM-Cre+ | Linear mixed effect model (LME) | Age | P16-P60 (32 mice, 80 recordings) | | | | Males: 46 recordings Females: 34 recordings | 0 | 1 |
| | | | Imaginary coherence (40-70 Hz) | 3 | Chi-Square: 15.024 | 0.001796 | | | |
| Figure 4A ii | Tukey's post-hoc test of LME-model | Condition | P16-17: PV-Cre+ - SOM-Cre+ | | z-value: 1.096 | 1 | | | |
| | | | P20-21: PV-Cre+ - SOM-Cre+ | | z-value: 4.380 | 0.000285 | | | |
| | | | P30-33: PV-Cre+ - SOM-Cre+ | | z-value: 2.008 | 0.714147 | | | |
| | | | P50-60: PV-Cre+ - SOM-Cre+ | | z-value: 4.431 | 0.000235 | | | |
| Figure 4A iii<br><br>PV-Cre+ | Linear mixed effect model (LME) | Age | P16-P60 (34 mice, 82 recordings) | | | | Males: 42 recordings Females: 40 recordings | 0 | 1 |
| | | | Imaginary coherence (20-35 Hz) | 3 | Chi-Square: 4.8528 | 0.1829 | | | |
| Figure 4A iii<br><br>SOM-Cre+ | Linear mixed effect model (LME) | Age | P16-P60 (32 mice, 81 recordings) | | | | Males: 45 recordings Females: 36 recordings | 0.0735 | 0.7864 |
| | | | Imaginary coherence (20-35 Hz) | 3 | Chi-Square: 2.4053 | 0.4926 | | | |
| Figure 4A iii | Tukey's post-hoc test of LME-model | Condition | P16-17: PV-Cre+ - SOM-Cre+ | | z-value: 2.006 | 1 | | | |
| | | | P20-21: PV-Cre+ - SOM-Cre+ | | z-value: 0.163 | 1 | | | |
| | | | P30-33: PV-Cre+ - SOM-Cre+ | | z-value: 0.184 | 1 | | | |
| | | | P50-60: PV-Cre+ - SOM-Cre+ | | z-value: 0.846 | 1 | | | |

| | | | | | | | | | |
|---|---|---|---|---|---|---|---|---|---|
| Figure 4A iv<br><br>PV-Cre+ | Linear mixed effect model (LME) | Age | P16-P60 (34 mice, 82 recordings) | | | | Males: 42 recordings Females: 40 recordings | 0 | 1 |
| | | | Imaginary coherence (40-70 Hz) | 3 | Chi-Square: 3.7179 | 0.2936 | | | |
| Figure 4A iv<br><br>SOM-Cre+ | Linear mixed effect model (LME) | Age | P16-P60 (32 mice, 81 recordings) | | | | Males: 45 recordings Females: 36 recordings | 0.0735 | 0.7864 |
| | | | Imaginary coherence (40-70 Hz) | 3 | Chi-Square: 8.9894 | 0.02943 | | | |
| Figure 4A iv | Tukey's post-hoc test of LME-model | Condition | P16-17: PV-Cre+ - SOM-Cre+ | | z-value: 0.737 | 1 | | | |
| | | | P20-21: PV-Cre+ - SOM-Cre+ | | z-value: 2.495 | 0.28996 | | | |
| | | | P30-33: PV-Cre+ - SOM-Cre+ | | z-value: 1.434 | 1 | | | |
| | | | P50-60: PV-Cre+ - SOM-Cre+ | | z-value: 2.437 | 0.32604 | | | |
| Figure 4B<br><br>(left) | Linear mixed effect model (LME) | Age | P16-P60 (53 mice, 109 recordings) | | | | Males: 53 recordings Females: 56 recordings | 0 | 1 |
| | | | Tiling coefficient (5ms) percentage change | 3 | Chi-Square: 10.378 | 0.01561 | | | |
| | | | P16-17: (12 mice, 363 unit pairs) | | | | | | |
| | | | P20-21: (13 mice, 1636 unit pairs ) | | | | | | |
| | | | P30-33: (15 mice, 591 unit pairs ) | | | | | | |
| | | | P50-60: (13 mice, 2350 unit pairs ) | | | | | | |
| | Tukey's post-hoc test of LME-model | Condition | P20-21 – P16-17 | | z-value: 1.691 | 0.3637 | | | |
| | | | P30-33 – P16-17 | | z-value: 0.116 | 0.9074 | | | |
| | | | P50-60 – P16-17 | | z-value: 2.677 | 0.0377 | | | |
| | | | P30-33 – P20-21 | | z-value: -1.657 | 0.3637 | | | |
| | | | P50-60 – P20-21 | | z-value: 1.089 | 0.5524 | | | |
| | | | P50-60 – P30-33 | | z-value: 2.732 | 0.0377 | | | |
| Figure 4B<br><br>(right) | Linear mixed effect | Age | P16-P60 (34 mice, 82 recordings) | | | | Males: 53 recordings Females: | 0 | 1 |

| | | | | | | | | | |
|---|---|---|---|---|---|---|---|---|---|
| | model (LME) | | Tiling coefficient (5ms) percentage change | 3 | Chi-Square: 2.968 | 0.3966 | 57 recordings | | |
| | | | P16-17: (12 mice, 744 unit pairs) | | | | | | |
| | | | P20-21: (13 mice, 887 unit pairs ) | | | | | | |
| | | | P30-33: (15 mice, 544 unit pairs ) | | | | | | |
| | | | P50-60: (13 mice, 1512 unit pairs ) | | | | | | |
| | Tukey's post-hoc test of LME-model | Condition | P20-21 – P16-17 | | z-value: 0.291 | 1 | | | |
| | | | P30-33 – P16-17 | | z-value: -0.650 | 1 | | | |
| | | | P50-60 – P16-17 | | z-value: 0.920 | 1 | | | |
| | | | P30-33 – P20-21 | | z-value: -0.956 | 1 | | | |
| | | | P50-60 – P20-21 | | z-value: 0.610 | 1 | | | |
| | | | P50-60 – P30-33 | | z-value: 1.668 | 0.571 | | | |
| Figure 4C (left) | Linear mixed effect model (LME) | Age | P16-P60 (53 mice, 110 recordings) | | | | Males: 61 recordings Females: 50 recordings | 0 | 1 |
| | | | Tiling coefficient (5ms) percentage change | 3 | Chi-Square: 7.7342 | 0.05184 | | | |
| | | | P16-17: (13 mice, 1771 unit pairs) | | | | | | |
| | | | P20-21: (13 mice, 14280 unit pairs ) | | | | | | |
| | | | P30-33: (8 mice, 4322 unit pairs ) | | | | | | |
| | | | P50-60: (15 mice, 8731 unit pairs ) | | | | | | |
| | Tukey's post-hoc test of LME-model | Condition | P20-21 – P16-17 | | z-value: 2.618 | 0.0531 | | | |
| | | | P30-33 – P16-17 | | z-value: 0.530 | 1 | | | |
| | | | P50-60 – P16-17 | | z-value: 1.170 | 0.7256 | | | |
| | | | P30-33 – P20-21 | | z-value: -2.026 | 0.2140 | | | |
| | | | P50-60 – P20-21 | | z-value: -1.560 | 0.4751 | | | |
| | | | P50-60 – P30-33 | | z-value: 0.589 | 1 | | | |
| Figure 4C (right) | Linear mixed effect | Age | P16-P60 (49 mice, 112 recordings) | | | | Males: 61 recordings Females: | 0 | 1 |

| | | | | | | | |
|---|---|---|---|---|---|---|---|
| | model (LME) | | Tiling coefficient (5ms) percentage change | 3 | Chi-Square: 3.1761 | 0.3653 | 51 recordings |
| | | | P16-17: (12 mice, 3339 unit pairs) | | | | |
| | | | P20-21: (13 mice, 2809 unit pairs ) | | | | |
| | | | P30-33: (15 mice, 4762 unit pairs ) | | | | |
| | | | P50-60: (13 mice, 8614 unit pairs ) | | | | |
| | Tukey's post-hoc test of LME-model | Condition | P20-21 – P16-17 | | z-value: 0.342 | 1 | |
| | | | P30-33 – P16-17 | | z-value: 1.088 | 1 | |
| | | | P50-60 – P16-17 | | z-value: 1.710 | 0.524 | |
| | | | P30-33 – P20-21 | | z-value: 0.703 | 1 | |
| | | | P50-60 – P20-21 | | z-value: 1.258 | 1 | |
| | | | P50-60 – P30-33 | | z-value: 0.469 | 1 | |

Legend Table S1: WT: wildtype, LME: linear mixed-effect model, GLME: generalized linear mixed-effect model, P: postnatal day, SDR: spectral dependency ratio, FS: fast-spiking, RS: regular-spiking, L: layer, CI: confidence interval, MI: modulation index, SUA: single-unit activity, gPDC: generalized partial directed coherence
